# Genetic integration with cell-specific nucleosome positioning resolves causal relationships underlying chromatin accessibility profiles

**DOI:** 10.1101/2025.09.09.674883

**Authors:** Xiaoou Wang, Catherine C. Robertson, Arushi Varshney, Nandini Manickam, Peter Orchard, Markku Laakso, Jaakko Tuomilehto, Timo A. Lakka, Karen L. Mohlke, Michael Boehnke, Laura J. Scott, Heikki A. Koistinen, Francis S. Collins, Stephen C.J. Parker

## Abstract

Cell type-specific chromatin accessibility QTL (caQTL) mapping is a promising approach to understand genetic control of chromatin landscapes and identify regulatory mechanisms underlying GWAS associations. However, current caQTL studies lack resolution and do not distinguish nucleosome-free regions (NFR) from positioned nucleosomes. Here, we leverage statistical modeling of fragment position and length to decompose ATAC-seq profiles into NFRs and phased nucleosomes. With single nucleus (sn)ATAC-seq from 281 human muscle biopsies, we map cell type-specific genetic effects on NFRs (76,027 nfrQTLs) and nucleosome occupancy (24,623 nucQTLs) across skeletal muscle cell types. Colocalization and causal inference between nucQTLs and nearby nfrQTLs and show that nfrQTLs are substantially more likely to causally influence nucQTLs and phase adjacent nucleosomes, indicating a causal relationship in shaping chromatin profiles. Hundreds of nfrQTLs colocalize with GWAS signals for muscle-related traits, including grip strength, atrial fibrillation, and fasting insulin, and the majority of colocalizing signals mapped to credible sets overlapping the corresponding nfrPeak. This approach adds mechanistic insights for how variants underlying caQTLs and GWAS signals exert their *cis* regulatory effects by initially modifying NFR accessibility and subsequently shaping broader chromatin landscapes, nucleosome positioning, gene expression, and ultimately higher-level traits and disease.

## Introduction

Genome-wide association studies (GWAS) have identified thousands of genetic associations with complex human diseases and traits^1^. Over 90% of GWAS signals map to noncoding regions and are hypothesized to function through gene regulatory mechanisms^2^. To nominate candidate genes, many studies integrate GWAS results with gene expression quantitative trait locus (eQTL) maps, leading to mechanistic hypotheses about how candidate variants influence disease risk through tissue-specific gene expression^3–9^. However, many GWAS associations do not colocalize with current eQTL maps^6,10^, limiting the power of this approach to resolve causal genes and mechanisms. The lack of eQTL colocalization at GWAS signals likely reflects several challenges: limited cell type resolution of current eQTL maps, which are primarily based on bulk tissue-level RNA-sequencing^5,11^; insufficient statistical power in eQTL studies^9^; failure to capture dynamic cell states that are critical to trait physiology^10^; and a mismatch in selective pressure between variants with large effects on gene regulation versus disease^12^. While some of these limitations are being addressed through eQTL studies with larger sample sizes^9,13^ and using single cell profiling^14–16^, evidence suggests that even large-scale eQTL studies with thousands of participants may remain underpowered to fully resolve disease-relevant regulatory mechanisms^9,12^.

Chromatin accessibility QTL (caQTL) mapping is a powerful, complementary approach for identifying molecular effects of trait-associated variants and refining credible sets at GWAS signals^17^. Relative to eQTLs, caQTL mapping is often more statistically powerful^18^. For example, paired cell type-specific caQTL and eQTL mapping in human muscle identified >10-fold more caQTLs than eQTLs across muscle cell types and more GWAS associations for muscle-related traits colocalized with caQTLs than eQTLs^19^, underscoring the promise of caQTL mapping for nominating regulatory mechanisms at GWAS signals. However, colocalizing caQTLs do not directly implicate causal genes at GWAS signals and, while they can point to candidate regulatory elements, the regions of accessibility prioritized by caQTLs can be broad and overlap multiple credible variants, making experimental validation challenging. New approaches are needed for pinpointing causal variants and regulatory regions at GWAS signals.

Most caQTL analyses rely on ATAC-seq, which quantifies chromatin accessibility by measuring susceptibility to Tn5 transposase integration^20^. ATAC-seq can be applied to a wide variety of human tissues and cell types^21^ and has been adapted to single nucleus platforms (snATAC-seq)^22,23^. Regions of high accessibility detected by ATAC-seq signal reflect a mix of regulatory events, including nucleosome displacement, transcription factor (TF) binding, and nucleosome positioning. However, standard bioinformatic processing of ATAC-seq data collapses these regulatory processes into one signal, ignoring valuable information present in the distribution and length of ATAC-seq fragments, and peak calling procedures (e.g., MACS2 narrowPeak^24^) tend to identify accessible regions encompassing both TF-bound sites and positioned nucleosomes surrounding the DNA-protein interaction.

Alternative approaches to analyzing ATAC-seq data can offer a more nuanced view of regulatory activity. For instance, TF footprint analysis uses patterns in ATAC-seq data to infer precise sites of TF binding^25^. However, this approach is limited by the brief residence time of most TFs on DNA, which often fails to produce a detectable footprint^26,27,28^. Another strategy is to use ATAC-seq read lengths to infer nucleosome-free regions (NFRs) and nucleosome positioning^29,30^, providing high-resolution insights into chromatin architecture genome-wide. Here, we use a new statistical ATAC-seq fragment partitioning approach and nucleosome fingerprints^30^ to decompose genetic control of chromatin dynamics into two components: accessibility in nucleosome-free regions (NFRs) (nfrQTLs) and nucleosome occupancy in regions with well-positioned nucleosomes (nucQTLs). Using snATAC-seq data from skeletal muscle cell types across 281 Finnish individuals^19^, we generate cell type-specific nfrQTL and nucQTL maps. We use causal inference to show that genetic control of chromatin accessibility landscapes is mediated through genetic effects on NFR accessibility, which in turn shapes adjacent nucleosome occupancy. We further show that nfrQTLs are more enriched for GWAS heritability than nucQTLs, and we provide empirical evidence supporting the corollary that GWAS colocalization with caQTLs is most often driven by causal variants in NFRs. Our work introduces a new framework for modeling genetic regulation of chromatin dynamics, enabling more precise identification of causal variants and mechanisms at GWAS loci and revealing causal relationships underlying genome-wide chromatin accessibility profiles.

## Results

### Calling NFRs and nucleosomes in five muscle cell types

We identified NFRs and nucleosome positioning in five muscle cell types by analyzing fragment length distributions and structured nucleosome fingerprints in a snATAC-seq dataset of skeletal muscle biopsies from 281 Finnish participants (**Fig 1a-b, Supplementary Fig 1**)^19,30^. First, for each participant, we extracted cell type-specific ATAC-seq profiles from the snATAC-seq data for the five most abundant cell types (type 1, type 2a, and type 2x muscle fibers; endothelial cells; and fibro-adipogenic progenitor (FAP) cells) using published clustering and cell type annotations^19^. Next, to map NFRs, we partitioned ATAC-seq data from each cell type within each participant into nucleosome-free and nucleosome-bound fragments. To identify an optimal threshold for classifying fragments as nucleosome-free versus nucleosome-bound in each cell type and participant, we applied multi-Otsu thresholding^31^, an image segmentation method, to the ATAC-seq fragment length distribution (**Fig 1a**). Empirically determined thresholds were consistent with a nucleosome core bound by 145-147 base pairs of DNA^32^ (mean multi-Otsu threshold for NFRs = 132.2 bp; standard deviation = 3.0 bp) and was consistent across individuals, cell types, and experimental batches (**Supplementary Fig 2a**). After partitioning ATAC-seq data according to participant- and cell type-specific thresholds, we called per-cell type NFR summits using nucleosome-free fragments merged across participants and expanded summits by 75bp on each side to define 151bp wide “nfrPeaks” (**Fig 1b**). For comparison, we also called summits on unpartitioned data and similarly extended summits by 75bp to define 151bp wide “caPeaks”^24^.

**Figure 1.**
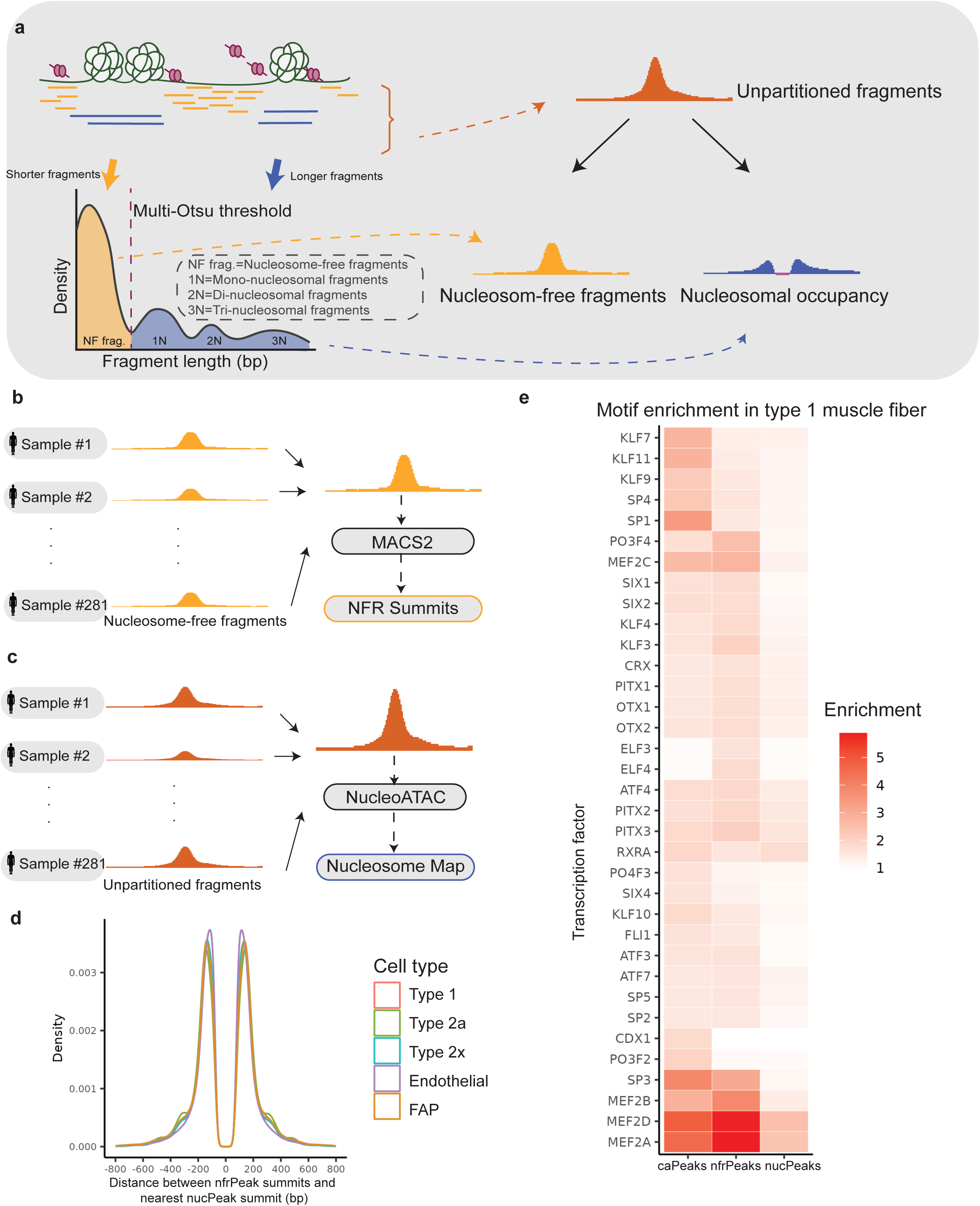
Decomposing chromatin accessibility into nucleosome-free regions (NFR) and nucleosome positioning. **a.** Multi-Otsu thresholding was applied to partition ATAC-seq data for each cell type in each participant into shorter nucleosome-free fragments and longer nucleosome-bound fragments. Nucleosome occupancy signal was quantified by subtracting nucleosome-free signal from nucleosome-bound signal. **b.** Merged nucleosome-free fragments were used to call NFR summits for each cell type. **c**. Unpartitioned fragments were used to call nucleosomes for each cell type. **d.** The distance between each nfrPeak and the closest nucPeak by cell type. **e.** Motif enrichment analyses on caPeaks, nfrPeaks, and nucPeaks in type 1 muscle fiber (showing motifs with enrichment ratio >1.5 in any peak category).

To map nucleosomes, we used NucleoATAC, which compares observed ATAC-seq fragment patterns to a “nucleosome fingerprint” - a characteristic density of fragment lengths in the 300bp region surrounding a nucleosome center - while normalizing for sequence bias and read depth^30^. We excluded nucleosomes with centers overlapping nfrPeaks and extended centers by 100 bp on each side to define 201bp wide “nucPeaks” (**Fig 1c**).

In each cell type, we called hundreds of thousands of peaks in each category (nfrPeaks, caPeaks, and nucPeaks) (**Supplementary Fig 2b**). To confirm that nfrPeaks and nucPeaks exhibited expected features, we investigated the positional relationship between nfrPeaks and nucPeaks and patterns in TF motif enrichment across caPeak, nfrPeak, and nucPeak regions. The median distance between nucPeaks and the nearest nfrPeak was 152bp, and 50% of nucPeak summits were located between 117 bp (25 percentile) and 206 bp (75 percentile) from the nearest nfrPeak summit (**Fig 1d**). Across all cell types, we found less pronounced TF motif enrichment patterns in nucPeaks compared to nfrPeaks and caPeaks (**Fig 1e, Supplementary Fig 3a-e**). We saw broad enrichment of TF motifs within nfrPeaks and caPeaks, with particularly striking motif enrichment for members of the MEF2 family (**Fig 1e, Supplementary Fig 3a-e**). In type 1, type 2a, type 2x, and FAP cells, we observed higher enrichment for TFs from the MEF2 family in nfrPeaks relative to caPeaks (**Fig 1e, Supplementary Fig 3a-c & e**). Together, these results are consistent with nfrPeaks mapping to sites of nucleosome displacement due to TF binding, nucPeaks corresponding to accessibility due to highly positioned nucleosomes, and caPeaks representing a mixture of NFRs and nucleosomes.

### Partitioned nfrQTL and nucQTL mapping identifies over a hundred thousand QTLs

We next investigated genetic control of NFRs and nucleosome positioning by mapping *cis* QTL effects on accessibility in nfrPeaks (nfrQTLs) and nucleosome occupancy in nucPeaks (nucQTLs) (**Fig 2a**). To build an NFR count matrix, we quantified nucleosome-free fragments overlapping nfrPeaks in each participant (**Fig 1a**, yellow tracks). To generate a nucleosome occupancy matrix, we first inferred nucleosome occupancy signal in each participant by subtracting smoothed nucleosome-free fragment signal from nucleosome-bound signal (Methods – Quantifying nucleosome occupancy). We then calculated average nucleosome occupancy across nucPeak regions in each participant to build a nucleosome occupancy score matrix for QTL analyses (**Fig 1a**, blue tracks).

**Figure 2.**
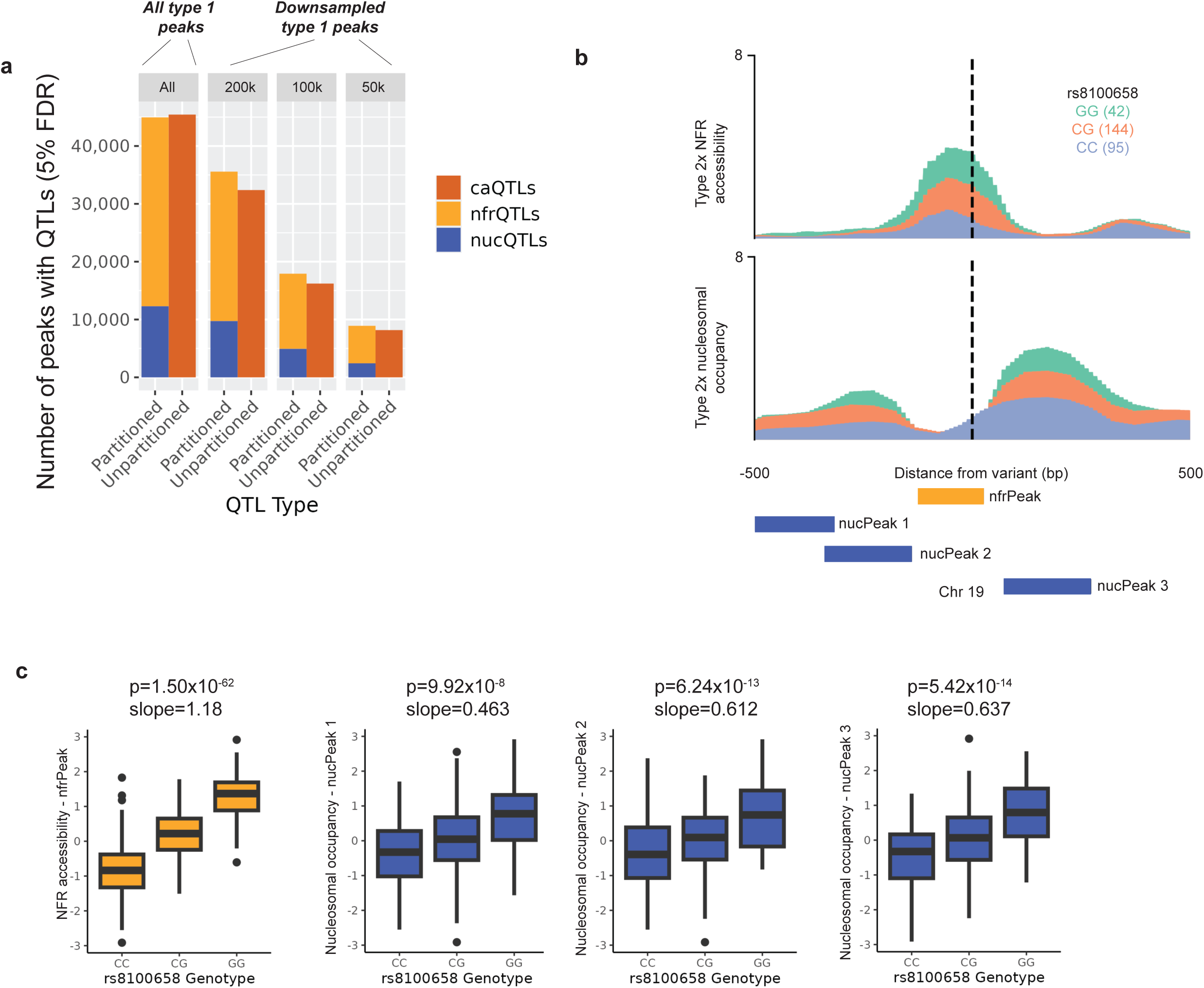
Discovery of *cis*-QTLs regulating nfrPeaks, nucPeaks, and caPeaks in five muscle cell types. **a.** The number of type 1 peaks with significant QTLs is reported for both unpartitioned (caQTL) and partitioned (nfrQTL and nucQTL) datasets. Results are categorized based on the downsampling experiment, which included all peaks and subsets of 200k, 100k, and 50k peaks. **b.** A layered plot showing NFR accessibility (top) and nucleosome occupancy (bottom) in type 2x muscle fibers surrounding variant rs8100658 on chromosome 19. Variant rs8100658 is located within nfrPeak chr19:34,994,613-34,994,764, which is associated with three nearby nucPeaks: nucPeak 1 (chr19:34,994,215-34,994,416), nucPeak 2 (chr19:34,994,396-34,994,597), and nucPeak 3 (chr19:34,994,812-34,995,013). **c.** Box plots display effect sizes and trends for the identified nfrPeak and nucPeaks.

We optimized *cis-*QTL models for each cell type and peak category by selecting observed and latent covariates that maximized power for QTL discovery (**Supplementary Fig 4a-b**). For observed covariates, we considered participant age, sex, and body mass index (BMI), experimental batch, number of nuclei, transcription start site (TSS) enrichment, and median fragment length (measured on chromosome 22). For latent variables, we considered genotype- and phenotype-derived principal components (PCs). Phenotype-derived PCs were calculated based on inverse rank normalized cell type-specific accessibility or occupancy matrices. Phenotypic PCs showed strong correlations with observed covariates, including sex, BMI, experimental batch, number of nuclei, and TSS enrichment (**Supplementary Fig 5-7**). We found that models incorporating genotype and phenotype PCs were more powerful than models that also included observed covariates (**Supplementary Fig 4a-b**). For downstream analyses, we proceeded with optimized models including the top five genotype PCs and the number of phenotype PCs (between 4-40) that yielded the greatest number of significant QTLs (**Supplementary Fig 4a-b, Supplementary Table 1**).

Our optimized QTL models identified 76,027 nfrQTLs and 24,623 nucQTLs across the five muscle cell types (FDR < 0.05 after two-step multiple testing correction) (**Fig 2a, Supplementary Fig 8&9**). We discovered the most nfrQTLs and nucQTLs in the most abundant muscle cell types— type 1, type 2a, and type 2x muscle fiber (**Fig 2a**, **Supplementary Fig 9**). We highlight a type 2x nfrQTL with three nearby nucQTLs on chromosome 19 (**Fig 2b-c**). At this locus, individuals with the rs8100658 alternative allele (G) showed higher chromatin accessibility in the nfrPeak and higher nucleosome occupancy in all three nucPeaks than those with the reference allele (C) (**Fig 2b-c**).

For comparison, we also mapped caQTLs using caPeak count matrices generated by quantifying unpartitioned fragments overlapping caPeaks in each participant. We normalized matrices and optimized models using the same approaches as used for nfrQTL and nucQTL mapping. We discovered slightly more caQTLs than nfrQTLs and nucQTLs combined (**Fig 2a, Supplementary Fig 8**). For example, in type 1 muscle fibers, we identified 45,433 peaks with significant caQTLs (FDR < 5%), while we identified 32,665 significant nfrQTL peaks and 12,288 significant nucQTL peaks (total of 44,953 peaks) (**Fig 2a**). However, this comparison is complicated by differing numbers of features included in the QTL scans and differing degrees of co-regulation among caPeaks as compared to nfrPeaks and nucPeaks. To address differing numbers of starting peaks, we performed downsampling experiments. When starting caQTL, nfrQTL, and nucQTL scans with the same number of peaks, the combined number of significant nfrQTLs and nucQTLs discovered was comparable to the number of significant caQTLs discovered (**Fig 2a, Supplementary Fig 8a & c**). To address co-regulation among peaks, we quantified the “effective” number of independent caPeaks or combined nfrPeaks and nucPeaks with QTLs using dimension reduction. Among peaks with significant QTLs, more peaks were required to explain 90% of the variance in accessibility profiles at partitioned nfrPeak and nucPeak regions than unpartitioned caPeak regions, suggesting more independent regulatory QTLs are identified with partitioning (**Supplementary Table 2**). Finally, to understand the extent to which nfrQTL and nucQTL analyses identify the same versus distinct regulatory regions as caQTLs, we analyzed the positional overlap between nfrPeaks, nucPeaks, and caPeaks with significant QTLs (**Supplementary Fig 9 & 10**). When we defined overlap as a minimum of 50% reciprocal overlap between peaks, the fraction of nfrQTL peaks that overlapped a caQTL peak ranged from 83.2-85.4% across cell types, and the fraction of nucQTL peaks that overlapped a caQTL peak was lower, ranging from 7.2-11.55% across cell types (Supplementary Fig 9). When we relaxed the criteria for overlap, requiring only 1bp of overlap between peaks, 88.1-93.3% of nfrQTLs and 82.5-92.2% of nucQTLs had peaks overlapping a caQTL peak.

### Over 70% of nucQTLs colocalize with nearby nfrQTLs

We next investigated evidence supporting shared causal variants underlying nfrQTLs and nearby nucQTLs (**Fig 3a**). We first fine mapped all significant nfrQTL and nucQTL associations using the Sum of Single Effects model (SuSiE) ^3^. Across all cell types, fine mapping converged to define credible sets for 40,189 out of 76,027 (52.9%) nfrQTLs and 12,886 out of 24,623 (52.3%) nucQTLs (**Supplementary Fig 11a-b**). Across cell types, 2,387 nfrQTL credible sets and 510 nucQTL credible sets contained just one variant (**Supplementary Fig 11a-b**). We then tested for formal colocalization between pairs of nfrQTLs and nucQTLs in close proximity (< 5 kb) using coloc v5 ^8^. Our colocalization analysis revealed 16,918 unique pairs of colocalized nfrQTLs and nucQTLs, which included 8,910 (22.2%) nfrQTL credible sets and 9,325 (72.4%) nucQTL credible sets across all five cell types (posterior probability, PPH4 > 0.8, **Supplementary Table 3**). To illustrate, we highlight the rs8100658 locus on chromosome 19 (as shown in **Fig 2b-c**), which has strong evidence of colocalization between the nfrQTL (nfrPeak: chr19:34,994,613-34,994,764) and nucQTL (nucPeak: chr19:34,994,812-34,995,013) signals in type 2x cells (PPH4>0.99) (**Fig 3b-c**). In this locus, rs8100658 is the lead variant for both the nfrQTL and the nucQTL (**Fig 3b-c**) and lies within the nfrPeak (chr19:34,994,611-34,994,762) (**Fig 2b**).

**Figure 3.**
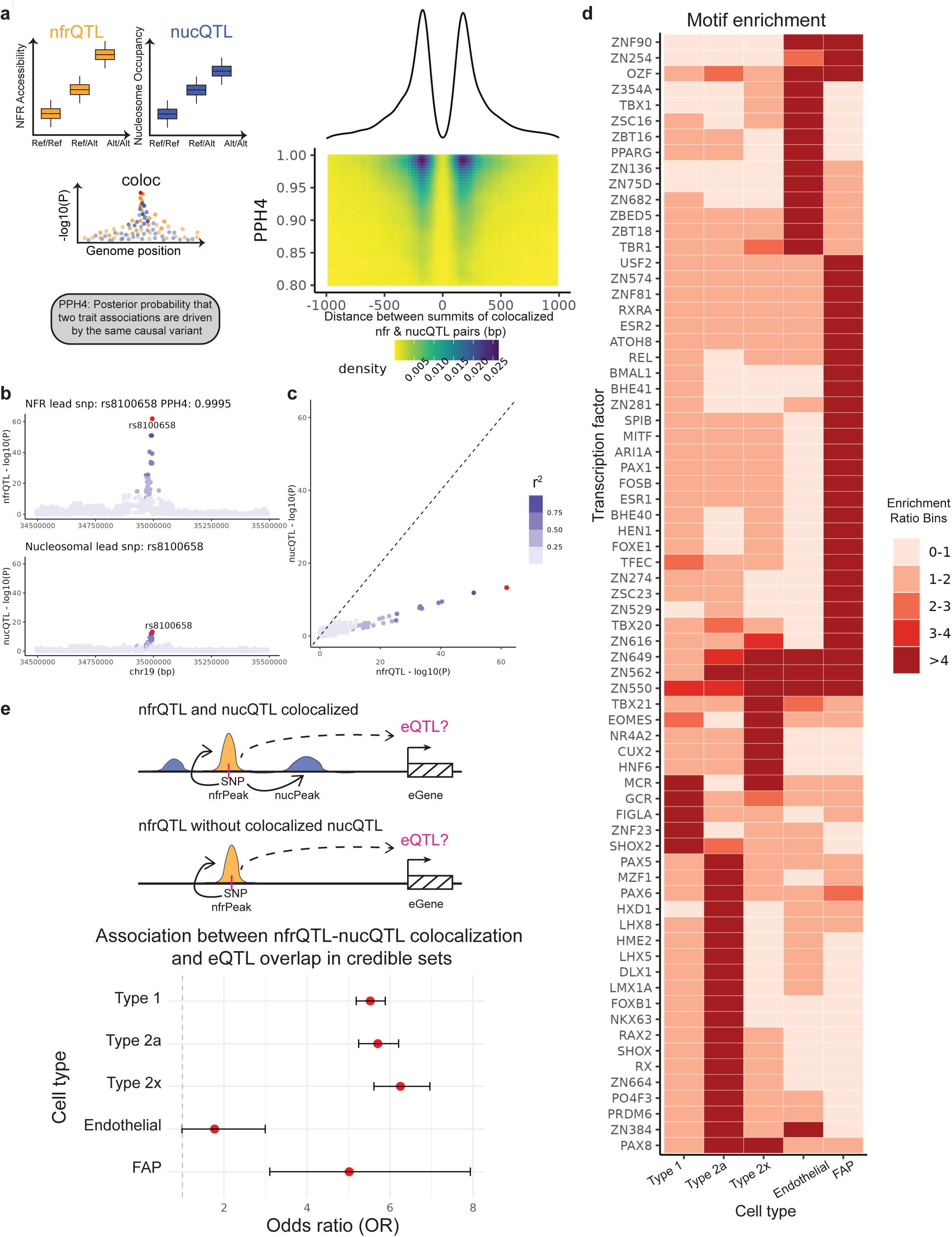
Colocalization patterns between nfrQTLs and nucQTLs. **a.** Left: a cartoon illustration of colocalization between an nfrQTL and a nucQTL. Right: the relationship between PPH4 and the distance between colocalized nfrQTL and nucQTL peak summits. **b.** Motif enrichment of nfrQTL peaks that colocalize with nucQTLs (PPH4 > 0.8) across five cell types. nfrQTLs without colocalized nucQTLs (PPH4 < 0.2) were used as control sequences. Showing motifs with enrichment ratio > 5 in any cell type. **c.** Locuszoom plots for nfrQTL and nucQTL associations. **d.** A Locuscompare plot showing a colocalized nfrQTL (nfrPeak: chr19:34,994,613-34,994,764) and nucQTL (nucPeak: chr19:34,994,812-34,995,013) in type 2x cells, with variant rs8100658 identified as a potential lead variant for both peaks. **e.** Top: a cartoon illustrating how we tested for whether nfrQTLs with nucQTL colocalization are more likely to be eQTLs. Bottom: Logistic regression results showing odds of an nfrQTL having an eQTL effect depending on whether it has a colocalized nucQTL or not. **f.** Motif enrichment analyses on nfrQTL peaks colocalized with nucQTLs (PPH4 > 0.8), with non-colocalized nfrQTL peaks (PPH4 < 0.2) as controls. The heatmap shows motifs passing enrichment > 5 in at least one cell type (rows = motifs; columns = cell types) colored by the range of enrichment values.

Out of 8,910 nfrQTL credible sets, 4,697 (52.7%) colocalized with only one nucQTL and 4,213 (47.3%) colocalized with more than one nucQTL. Meanwhile out of 9,325 nucQTL credible sets, 5,175 (55.5%) colocalized with only one nfrQTL (**Supplementary Table 3**) and 4,150 (44.5%) with more than one nfrQTL. While we tested for colocalization for all nucQTL peaks within 5 kb of a nfrQTL peak, nucQTL peaks were typically within 500bp of their colocalizing nfrQTL peak, with a modal summit-to-summit distance of 167bp (**Fig 3a, Supplementary Fig 11e**). Among nfrQTL peaks colocalizing with more than one nucQTL peak, the distribution of summit-to-summit distances was multimodal (mode 1 = 166 bp; mode 2 = 311 bp) (**Supplementary Fig 11c & f**), implying these include nfrPeaks flanked by multiple phased nucleosomes.

Across cell types, distinct sets of motifs were enriched in nfrQTL peaks with colocalizing nucQTLs compared to those without colocalized nucQTLs (**Fig 3d**). For example, in type 2a muscle fiber, motifs for PAX5 (p=2.9 × 10^−5^) and PAX6 (p=1.1 × 10^−4^) were enriched in nfrPeaks associated with nfrQTL-nucQTL colocalizing pairs. PAX transcription factors are known pioneer factors capable of initiating chromatin remodeling^33^ and with some family members having established roles in myogenesis^34^. To explore the relationship between nucQTL colocalization and gene expression, we examined the overlap between nfrQTLs and cell type-specific eQTLs identified from snRNA-seq of the same 281 participants^19^. We first categorized nfrQTLs into two bins based on whether they had colocalized nucQTLs (**Fig 3e**, top). Then, we further categorized nfrQTLs based on whether the nfrQTL credible set overlapped an eQTL credible set defined in Varshney et al. 2024^19^ (**Fig 3e**, top). Using a logistic regression model adjusting for mean accessibility in the nfrPeak, we found that nfrQTLs with colocalized nucQTLs were more likely to overlap eQTLs in type 1 (OR = 5.2; p < 2.2 × 10^−16^), type 2a (OR = 5.2; p < 2.2 × 10^−16^), type 2x (OR = 5.6; p < 2.2 × 10^−16^), and FAP (OR = 3.1; p = 1.3 × 10^−11^) cells (**Fig 3e**, bottom). In summary, these analyses demonstrated that nfrQTL variants with effects on nucleosome occupancy are enriched for TF motifs in a cell type-specific manor, including for pioneer TFs, and are more likely to have effects on gene expression.

### Causal variants are more likely to overlap nfrPeaks than nucPeaks

We assessed the fraction of colocalized nfrQTL and nucQTL credible variants falling inside their corresponding nfrPeak or nucPeak. Across the QTL fine mapping posterior inclusion probability (PIP) spectrum, nfrQTL credible variants were more likely to overlap their respective nfrPeaks than nucQTL credible variants were to overlap their respective nucPeaks, and the difference was most pronounced among credible variants with higher PIPs (**Fig 4a**). In type 1 muscle fiber, nfrQTL credible set variants with PIP greater than 0.9 had a 42.5% chance of falling within the nfrPeak. In contrast, type 1 fiber nucQTL credible variants with PIP greater than 0.9 had only a 3.1% chance of being located within the nucPeak.

**Figure 4.**
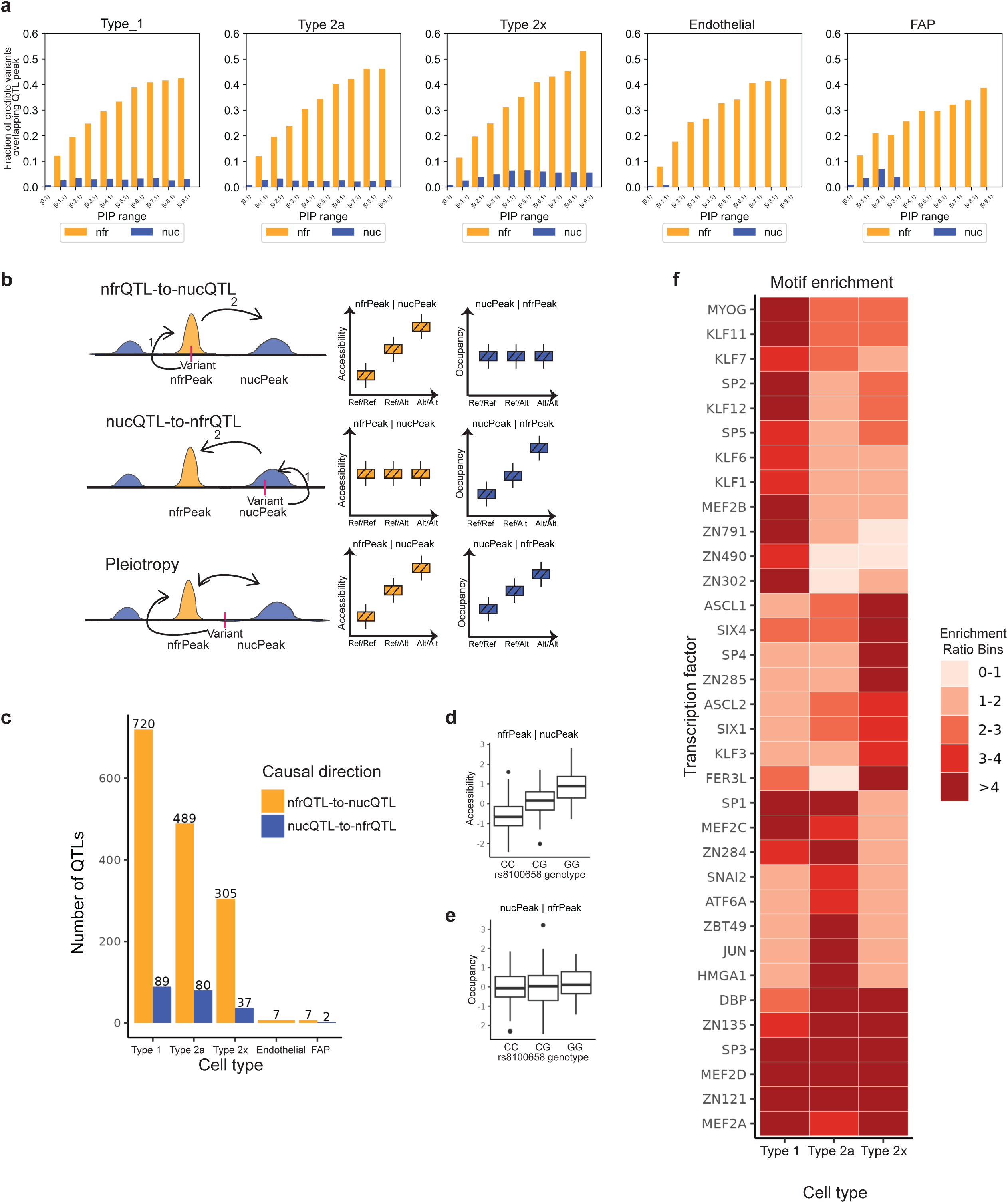
Causal relationships between nfrQTLs and nucQTLs. **a.** The probability of QTL credible variants being located within the nfrPeak or nucPeak for each cell type, evaluated across various PIP thresholds. By examining these probabilities, we assess the extent to which variants are spatially associated with open chromatin regions (nfrPeaks) or nucleosome positions (nucPeaks). **b.** A diagram outlines the three hypotheses assessed in the causal inference test. These are illustrated by regressing out either the nucleosome occupancy of the nucPeak for the instrumental variant (nfr|nuc) or the nfrPeak accessibility (nuc|nfr). Conditional QTL effects at three variants. X-axis is genotype dosage for each variant. Y-axis is conditional NFR accessibility or nucleosome occupancy signal - “nfr|nuc” is nfrPeak accessibility after conditioning on nucPeak occupancy (left) and “nuc|nfr” is nucPeak occupancy after conditioning on nfrPeak accessibility (right). **c.** Bar plots display the number of colocalized nfrQTL and nucQTL pairs with specified directionalities for each cell type. An example of nfr-to-nuc directionality in type 2x cells at rs8100658, when **d**. nfrPeak accessibility conditioned on nucPeak occupancy and **e.** nucPeak occupancy conditioned on nfrPeak accessibility. **f.** Selected motifs’ enrichment ratio heatmaps on nfrPeaks with causal directionality (nfrQTL-to-nucQTL). Heatmap shows enrichment ratios in 34 motifs with FDR<0.05 and enrichment ratio > 3 in any of the three cell types. Results in Endothelilal and FAP cells were not shown due to limited number of input sequences. PIP, posterior inclusion probability.

We next evaluated causal relationships between all colocalizing pairs of nfrQTLs and nucQTLs using a formal causal inference test (CIT)^35^ (**Fig 4b**). Across the five muscle cell types, we assigned causal directionality to 1,736 colocalizing pairs, with 1,528 pairs (88%) showing a nfrQTL-to-nucQTL direction and 208 pairs exhibiting a nucQTL-to-nfrQTL direction (**Fig 4c**). We highlight conditional QTL effects at the rs8100658 locus (as shown in **Fig 2b-c** and **Fig 3c-d**), which demonstrated nfrQTL-to-nucQTL causality (p=1.5×10^−4^) (**Fig 4d**). The strong effect of rs8100658 on nfrPeak accessibility is reflected even after conditioning on nucPeak occupancy (**Fig 4d**), while rs8100658 no longer influences nucPeak occupancy after conditioning on nfrPeak accessibility (**Fig 4e**).

In muscle fiber cell types (type 1, type 2a, and type 2x), nfrPeaks with causal effects on nucQTLs were enriched for binding motifs for the MEF2 family of transcription factors (**Fig 4f**). Enrichment was strongest in type 1 fibers (MEF2A enrichment ratio = 36, p = 2.15×10^−21^; MEF2D enrichment ratio = 36, p = 2.15×10^−21^) and also significant in type 2a (MEF2A enrichment ratio = 3.41, p = 4.1 × 10^−14^; MEF2D enrichment ratio = 4.42, p = 6.1 × 10^−16^) and type 2x fiber (MEF2A enrichment ratio = 4.92, p = 3.9×10^−10^; MEF2D enrichment ratio = 6, p = 1.9×10^−10^).

Together, the predominance of nfrQTL credible variants overlapping nfrPeaks and nfrQTL-to-nucQTL directionality among colocalizing nfrQTLs and nucQTLs implies that, most often, genetic variants first exert their effects in NFRs, which then influences positioning of adjacent nucleosomes. Further motif analyses indicate that, in muscle fiber, NFR-mediated genetic regulation of chromatin landscapes is partially mediated through altered binding of MEF2 family transcription factors.

### GWAS integration with nfrQTLs nominates molecular mechanisms

We tested GWAS enrichment for 384 traits in nfrQTL and nucQTL peaks (**Supplementary Table 4**)^19,36^ using linkage disequilibrium score (LDSC) regression-based enrichment analyses (**Fig 5a, Supplementary Fig 12**). Among traits that were significantly enriched in nfrQTL or nucQTL peaks (FDR<5%), the overwhelmingly majority showed more significant enrichment in nfrQTL peaks than nucQTL peaks (62 of 66 traits in type 1; 87 of 88 traits in type 2a; 79 of 79 traits in type 2x) (**Fig 5a**, **Supplementary Fig 12**). Focusing on traits related to muscle function that were enriched in nfrQTL peaks, including grip strength (UKBB, 46^36^), standing height (UKBB, 50^36^), basal metabolic rate (UKBB, 23105^36^), creatinine (UKBB, 30700^36^), type 2 diabetes (T2D)^37^, fasting insulin^38^, and atrial fibrillation^39^ (**Fig 5b, Supplementary Fig 13**), we used formal colocalization analysis to identify nfrQTLs colocalizing with GWAS signals (PPH4>0.8) (**Table 1**). Between 15% and 28% of GWAS signals colocalized with at least one nfrQTL (**Table 1**). Moreover, 9% to 20% of GWAS signals colocalized with a nfrQTL where a credible variant overlapped the nfrPeak (**Table 1**). Notably, out of the seven muscle-related traits considered, atrial fibrillation showed the highest GWAS enrichment in nfrQTL peaks (**Fig 5b, Supplementary Fig 13**) and had the highest frequency of credible sets colocalizing with nfrQTLs (28%). Among the 45 atrial fibrillation signals colocalizing with a nfrQTL, 33 contained credible variants overlapping the nfrPeak.

**Figure 5.**
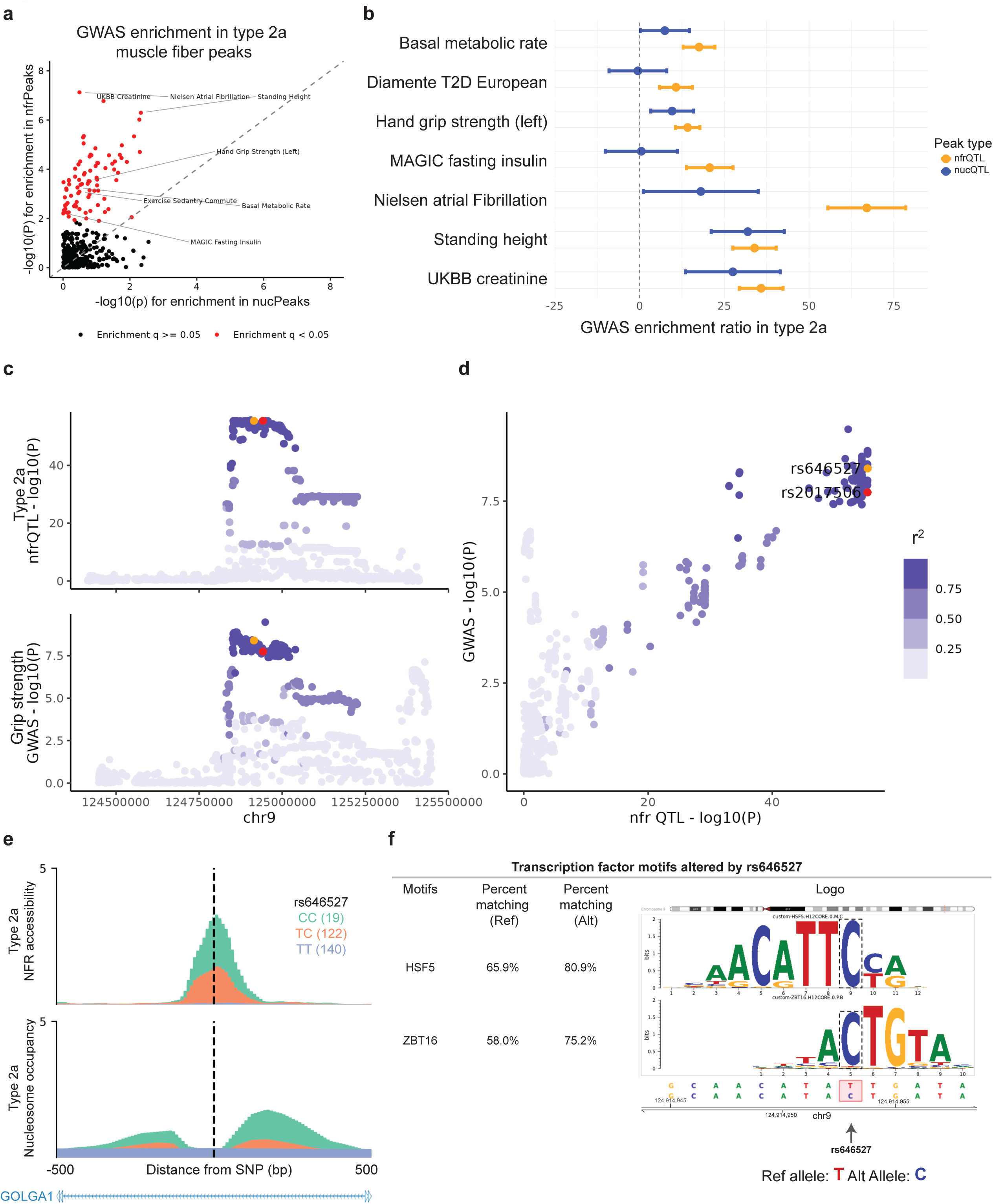
Colocalization between nfrQTLs and muscle-related GWAS traits. **a.** LDSC single enrichment analysis was conducted on 384 GWAS traits for Type 2a cells in nfrPeaks and nucPeaks with significant QTLs. Traits with enrichment q-values < 0.05 are highlighted in red, while those with insignificant enrichment are marked in black. The dashed line represents y=x. **b.** Seven muscle and metabolic-related GWAS traits were selected for fine mapping, and their enrichment values were compared between nfrPeaks and nucPeaks. Error bars represent enrichment standard errors. **c.** Locuszoom and **d.** Locuscompare plots illustrate the colocalization between a Type 2a nfrQTL (chr9:124,914,903-124,915,054) and the grip strength GWAS summary statistics (UKBB 50). Variants are colored by the r² value calculated from linkage disequilibrium (LD) matrices, with the potential lead variant, rs10986471, shown in red. Variant rs646527 shown as orange is within the credible set of rs10986471 and also lies within the Type 2a nrPeak (chr9:124,914,903-124,915,054) **e.** Layered plots show NFR accessibility (chr9:124,914,903-124,915,054) and nucleosome occupancy (chr9:124,914,679-124,914,880, chr9:124,915,036-124,915,237, and chr9:124,915,188-124,915,389) for this colocalization event in Type 2a cells. **f.** Predicted effects of rs646527 C allele on transcription factor binding motifs. Results were filtered to the motifs with allele effect size > 0.12 and p<0.005. The variant’s position within the motifs were boxed by the dashed lines.

**Table 1.**
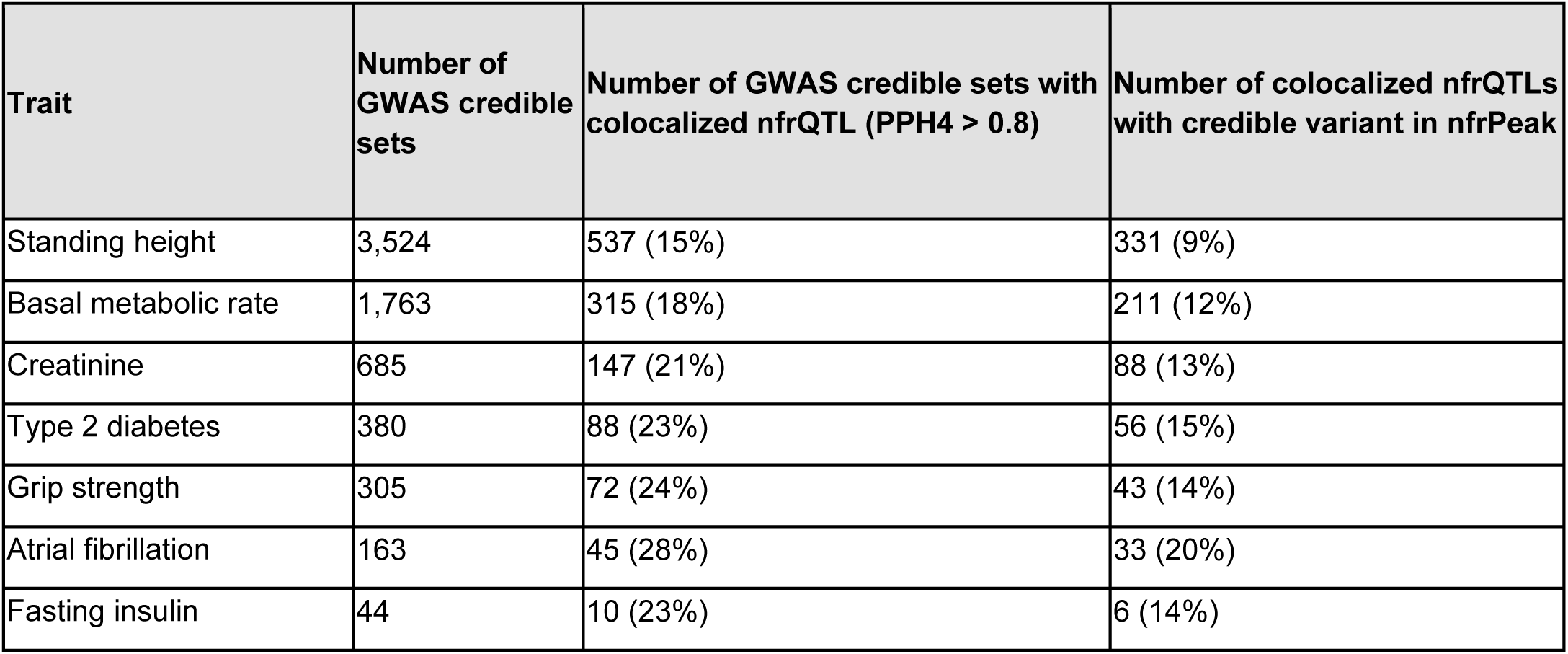
Summary of colocalization between nfrQTLs and GWAS signals.

To understand whether partitioned nfrQTL and nucQTL analyses identified distinct mechanisms from unpartitioned caQTL analyses, we compared grip strength GWAS signal colocalization with QTLs in type 2a muscle fiber across the three QTL categories (caQTLs, nfrQTLs, and nucQTLs). Out of 305 known grip strength GWAS signals, 55 colocalized with at least one QTL type (**Supplementary Fig 14**, PPH4>0.8). Most grip strength signals colocalizing with nfrQTLs and/or nucQTLs also colocalized with caQTLs (29/41). However, 12 signals colocalizing with partitioned nfrQTLs and/or nucQTLs did not colocalize with unpartitioned caQTLs; while 14 signals colocalizing with unpartitioned caQTLs did not colocalize with either a nfrQTL or a nucQTL.

On chromosome 9, we identified colocalization between a type 2a nfrQTL (nfrPeak region: chr9:124,914,903-124,915,054; lead variant rs2017506) with a grip strength association (PPH4=0.96) (**Fig 5c-f**). Within the colocalizing nfrQTL credible set, rs646527 overlapped the nfrQTL peak (**Fig 5e**). This nfrQTL also colocalized with three nucQTLs (**Fig 5e**), which themselves colocalized with the nfrQTL. Relative to the reference sequence (containing the T allele), the C allele at rs646527 increases accessibility in the nfrPeak and increases sequence similarity to binding motifs for the transcription factors HSF5 (80.9% versus 65.9% match) and ZBT16 (75.2% versus 58.0% match) (**Fig 5f**). In this locus, rs646527 represents a compelling candidate causal variant for grip strength, where the C allele is hypothesized to impair grip strength by increasing binding of one or more transcription factors in the chr9:124,914,903-124,915,054 region (nfrPeak) in type 2a fibers (PPH4=0.960). This example demonstrates the value of nfrQTL mapping for generating testable hypotheses about causal cell types and regulatory regions underlying GWAS traits.

## Discussion

Cell type-specific caQTL mapping is a promising approach to nominate regulatory mechanisms underlying GWAS associations. However, standard ATAC-seq analyses conflate accessibility induced by nucleosome phasing with accessibility due to TF binding and nucleosome displacement, leading to prioritization of broad regulatory element regions (caPeaks) with limited interpretability in terms of their molecular context. In this study, we introduced a new approach to mapping genetic effects on chromatin dynamics that distinguishes TF-bound regions from phased nucleosomes.

After decomposing snATAC-seq signal into cell type-specific NFR accessibility and nucleosome occupancy, we identified tens of thousands of nfrQTLs and nucQTLs in five muscle cell types. Through fine mapping, we found that causal variants for nfrQTLs are likely to overlap their corresponding nfrPeak, while causal variants for nucQTLs rarely overlap nucPeaks. Moreover, formal colocalization analyses demonstrated that most nucQTLs likely share a causal variant with a nearby nfrQTL. Causal inference testing supported the interpretation that, for the majority (>88%) of colocalizing nucQTL-nfrQTL pairs with causal directions, nfrQTLs play a causal role in regulating adjacent nucleosome occupancy. Our findings suggest that genetic association with nucleosome positioning is usually secondary to NFR changes. However, causal inference did detect a subset of colocalizing nfrQTLs and nucQTLs with pleiotropic effects or nucQTL-to-nfrQTL directionality, implying that some nucQTLs may reflect genetic control of nucleosome positioning through mechanisms independent of adjacent NFR regions and a limited set of NFR regions are themselves sensitive to such effects on nucleosomes^40^.

When we integrated nfrQTL mapping with GWAS associations for seven muscle-related traits, 15-28% of GWAS associations showed colocalization with one or more nfrQTLs. Among GWAS associations colocalizing with an nfrQTL, the majority had at least one credible variable overlapping a prioritized nfrPeak. These variants and the nfrPeaks they overlap implicate testable regulatory hypotheses, as demonstrated at rs646527, a non-coding variant which lies in an intronic enhancer in *GOLGA1*. The rs646527 alternative allele (C) increases nfrPeak accessibility, decreases grip strength, and disrupts motifs for multiple TFs, including ZBT16/ZBTB16 and the heat shock transcription factor HSF5. ZBTB16, also known as PLZF, is an essential regulator of muscle development and muscle-specific gene expression. It binds preferentially to accessible chromatin regions, and recent studies have shown that ZBTB16 localizes with other key transcription factors and chromatin regulators to help establish a chromatin interaction network in progenitor cells^41^. In the context of rs646527, the C allele increases NFR accessibility and enhances ZBTB16 binding, potentially driving dysregulated gene expression programs that impair muscle function, which manifests in decreased grip strength. This example illustrates how mapping nfrQTLs can facilitate prioritization of causal variants, mechanisms, and *cis-*regulatory elements (CREs) at GWAS loci.

In summary, our findings demonstrate that NFR and nucleosome inference, nfrQTL and nucQTL mapping, and nfrQTL colocalization with GWAS trait associations is a novel and effective strategy for prioritizing causal variants and regulatory elements with power and precision. This enhanced approach can empower functional follow-up of candidate causal variants using experimental approaches, which are costly and time-consuming. Finally, dissecting chromatin QTLs into NFR and nucleosome components reveals causal relationships underlying cell type-specific chromatin dynamics, providing insight in causal mechanisms underlying cell type-specific gene regulation.

## Methods

### Sample collection and data generation

The biological samples included in this study were sourced from 281 *vastus lateralis* muscle biopsies collected and characterized in the Finland-United States Investigation of NIDDM Genetics (FUSION study)^19^. We have provided full details of the sample collection protocols, including the ethical approval process, informed consent, and the specific criteria for sample inclusion and exclusion previously^19^. Briefly, all participants are of European ancestry and at least 18 years old. We obtained access to the samples through the FUSION collaboration, ensuring compliance with all regulations and agreements set forth by the original investigators and institutions. Processing of snATAC-seq and genotyping data was performed as described previously^19^. Briefly, snATAC-seq libraries were pre-processed using 10x Genomics Cell Ranger ATAC (v.1.0), nuclei were filtered using rigorous quality control steps^19^, and cell types were determined through joint clustering with the snRNA modality using Liger^42^. Using published clustering results^19^, cell type- and participant-specific ATAC-seq bam files were generated for the five most abundant cell types (type 1, type 2a, and type 2x muscle fibers; endothelial cells; and FAP cells).

### Partitioning ATAC-seq fragments using multi-Otsu thresholding

For each sample and each cell type, we used the SAMtools’ *view* function to isolate fragments from chromosome 22: *-bo sample_chr22.bam sample.bam chr22*. Considering only the properly mapped paired reads, we generated a list of insert sizes for chromosome 22 using SAMtools’ *view* function: *-f66 sample_chr22.bam | cut -f9 | awk ‘{print sqrt($0^2)}’ > sizes.* Next, we applied multi-Otsu statistical thresholding^31^ to the distribution of these insert sizes, selecting three thresholds which divided insert sizes into four classes: nucleosome-free, mono-nucleosomal (1N), di-nucleosomal (2N), and tri-nucleosomal (3N) fragments (**Fig 1b**). We then split each bam file into two bam files containing shorter and longer fragments, using the following SAMtools commands, where *t* is the first Otsu threshold: *view -h sample.bam | awk -v t ‘(substr($0,1,1)==“@“||($9>= -t && $9<=t))’ | view -b > shorter.bam* and *view -h sample.bam | awk -v t ‘(substr($0,1,1)==“@“||($9<= -t && $9>=t))’ | view -b > longer.bam*.

### Calling peaks from unpartitioned and nucleosome-free ATAC-seq reads

To define general “caPeaks” on unpartitioned ATAC-seq data, we called peaks using MACS2 (v2.2.9.1)^24^ with unpartitioned ATAC-seq bam files merged across samples for each cell type using the following command: *macs2 callpeak -t celltype.bed --outdir celltype_ca --bdg --SPMR -f BED -n celltype_ca -g hs --shift -50 --extsize 100 --seed 762873 --keep-dup all --nomodel --call-summit*. We extended inferred summits by 75 bp on each side to define the set of caPeaks.

To define NFR peak regions, we used the *shorter.bam* files (see previous section, “Partitioning ATAC-seq fragments using multi-Otsu thresholding”) merged across samples for each cell type. We merged *shorter.bam* files across samples and sorted the merged file using the SAMtools *merge* and *sort* functions, respectively. We then converted the merged per-cell type nucleosome-free bam files to bed files using BEDTools’ *bamtobed* function^43^ and called peaks using MACS2 (v2.2.9.1)^24^: *macs2 callpeak -t celltype_nfr.bed --outdir celltype_nfr -- bdg --SPMR -f BED -n celltype_nfr -g hs --shift -50 --extsize 100 --seed 762873 --keep-dup all --nomodel --call- summit.* We extended summits by 75 bp on both sides to define nfrPeaks for each cell type cluster.

### Calling nucleosome positions

We called nucleosome positions for each cell type using NucleoATAC (v0.3.4, GCC 8.5.0 20210514) ^30^ (**Fig 1c**). To run NucleoATAC, we first converted unpartitioned per-cell type bam files to bed files using BEDTools’ *bamtobed* function and generated broad peaks using MACS2 (v2.2.9.1)^24,43^ with the following command: *macs2 callpeak -t celltype.bed --outdir celltype --broad --bdg --SPMR -f BED -n celltype -g hs --shift -100 --extsize 200 --seed 762873 --keep-dup all --nomodel.* We then used the *run* command from NucleoATAC to call nucleosome positions: *run --bed celltype.broadPeak --bam celltype.bam --fasta hg38.fasta --out celltype_nuc.* We aligned reads to the GRCh38 reference assembly. To generate the most comprehensive map of nucleosome positions, we used the *nucmap_combined.bed* file, which combined both lower and higher resolution nucleosome calls from the *occ* and *nuc* functions. To avoid overlaps with the NFR, we used BEDTools’ *subtract* function^43^ to filter the inferred nucleosome positions that lay outside the NFR peaks and extended 100 bp on each side of the nucleosome positions to define nucleosome peaks for each cell type. To assess the distance and patterns between nfrPeaks and nucPeaks, we checked the distance between nfr summits and the closest nuc summit using BEDTools’ *closest* tool^43^, *-a nuc_summits.bed -b nfr_summits.bed -D ref.* We plotted the density for each cell type’s distance distribution in a +-800 bp window and identified the positionwith the highest density value was selected.

### Quantifying accessibility at NFR peaks and caPeaks

To quantify accessibility in caPeaks and nfrPeaks, we used the per-sample per-cell type unpartitioned bam and *shorter.bam* files, respectively. We used the *featureCounts* function (v2.0.6) ^44^ to count the number of mapped reads per feature, with the command: *featureCounts -F SAF -O --minOverlap 1 -T 16 -p -a celltype_nfr.saf -- donotsort --countReadPairs -o celltype_nfr_counts nfr*.bam.* The *.saf* file was converted from the *.bed* file, which included the defined caPeaks or nfrPeaks.

### Quantifying nucleosome occupancy

We created a separate BigWig file for each sample to capture nucleosome occupancy for the nucleosome positions of interest. After thresholding and splitting, we used the *bamCoverage* function (v3.5.4.post1) from deepTools ^45^ to generate coverage tracks for each sample’s *nfr.bam* and *longer.bam* files with the following commands:

bamCoverage --normalizeUsing BPM --blackListFileName ENCODE_blacklist --extendReads –binSize
*10 --smoothLength 30 --outFileName nfr.bw*
*bamCoverage --normalizeUsing BPM --blackListFileName ENCODE_blacklist --extendReads –binSize*
*50 --smoothLength 150 --outFileName longer.bw*

The blacklist from the ENCODE project was used to remove unstructured and problematic regions in the human genome ^46^. Since the *longer.bw* will include fragment pileup from nucleosome-bound fragments which extend into adjacent NFRs, we subtracted the *nfr.bw* fragment pileup signal from the *longer.bw* signal to quantify nucleosome occupancy (*nuc.bw*). We ran the subtraction using the *bigwigCompare* function (v3.5.4.post1): *bigwigCompare -b1 longer.bw -b2 nfr.bw --operation subtract --skipNAs --blackListFileName ENCODE_blacklist --outFileName nuc.bdg --outFileFormat bedgraph.* Since the subtraction introduced negative values, we used MACS2’s *bdgopt* (v2.2.9.1) ^24^ function to replace all negative values by 0 and converted the bedgraph file back to the BigWig format. Using the coverage information from *nuc.bw*, we built the nucleosome occupancy score matrix with the *bigWigAverageOverBed* function (v3.5.4.post1) from deepTools ^45^ to get the average score over the nucleosome peaks.

### Matrix filtering and normalization

For both the caPeak count matrices and the nfrPeak count matrices, we filtered for peaks with more than three reads in at least 5% of samples. For the nucleosome occupancy matrices, we filtered for nucleosome regions with an average nucleosome occupancy score >0.5 in at least 5% of samples. We normalized each matrix for sequencing depth using the Trimmed Mean of M (TMM) component normalization process to compute counts per million (CPM), as implemented in edgeR. We then inverse-normalized counts per million (CPM) values for each feature across samples.

### QTL mapping

For both caQTL, nfrQTL, and nucQTL scans, we used QTLtools (v1.3.1-25-g6e49f85f20)^47^ to perform cis-QTL scans over a *cis* window of 10 kb. We applied a two-step multiple testing correction: first, empirical p-values were obtained from permutations for each molecular phenotype; second, these p-values were adjusted across all tests using the Storey q-value method to control the false discovery rate (FDR). The scans included 1,000 permutations and were corrected for a FDR of less than 5%. We included the top five genotype principal components (PCs) as covariates in the analyses. Additionally, phenotype PCs, derived from the normalized count matrices, were incorporated, and the PCs were calculated with the sklearn.decomposition function from the scikit-learn package (v1.4.0). We conducted a systematic parameter sweep to explore the number of phenotype PCs and covariates in the QTL analysis model (**Supplementary Fig 4**). Meanwhile, we conducted another rounds of QTL scan to get nominal p-values between the nfr/nucPeaks and the genotypes. We ran the scan using the parameters *–nominal 1.0 –window 500000*.

### Downsampling peaks to compare caQTL, nfrQTL, and nucQTL discovery

To compare discovery power of caQTL, nfrQTL, and nucQTL mapping while accounting to for the overall number of features included in the analyses, we conducted a downsampling experiment, which started with the same number of caPeaks, nfrPeaks, and nucPeaks. When starting with 200k of randomly selected peaks, in type 1, we found 32,383 caPeaks with significant caQTLs, 25,864 significant nfrQTLs, and 9,725 significant nucQTLs (**Fig 2a**). In this case, when adding the number of peaks with significant nfrQTLs together with that of nucQTLs, we found more peaks than with caQTLs alone (**Fig 2a**). We observed a similar trend when randomly downsampling to 100k and 50k features. We also found the same trends not only in type 1 but also in the other four cell types that we studied (**Supplementary Fig 8**).

### Estimating the effective number of independent QTLs in caQTL versus nfrQTL and nucQTL discovery

To compare QTL discovery using nfrQTL and nucQTL mapping with discovery using caQTL mapping while accounting for co-regulation between peaks, we sought to estimate the effective number of independent caPeaks with caQTLs or independent nfrPeaks and nucPeaks with nfrQTLs or nucQTLs. We first constructed two peak matrices for each cell type: (1) caPeaks with significant caQTLs; (2) merged matrix of nfrPeaks with nfrQTLs and nucPeaks with nucQTLs. Next, we centered and scaled the matrices and ran PCA analysis using the prcomp function in the stats R package. We then approximated the number of independent columns within the QTL-associated peak matrices by estimating the number of PCs required to explain 90% of the variance in the matrix.

### QTL peaks overlapping

We used the BEDtools *intersect* function^43^ to reciprocally check the overlap between caQTL peaks and nfrQTL or nucQTL peaks. We defined overlap between two peaks using two criteria: (1) requiring at least 50% reciprocal overlap, meaning at least 50% of both peaks overlaps the other peak. This was implemented using the following option: *-wa -a {peakA} -b {peakB} -f 0.50 -r*; (2) requiring only 1bp overlap. This was implemented with the following option: *-wa -a {<u>peakA</u>} -b {peakB}*.

### QTL fine mapping and colocalization

We used the SUm of SIngle Effects (SuSiE) model^48^ to build 95% credible sets for significant nfrQTLs and nucQTLs. We used the corrected normalized count and score matrices and the genotype dosage information for each variant as inputs to run the SuSiE model. The phenotypic data were normalized and corrected for covariates by QTLtools’ *correct* function (v1.3.1-25-g6e49f85f20) ^47^. We checked the variants that are within 500kb window size to the nfr and nucQTLs and built the 95% credible sets by setting: *L = 10, estimate_residual_variance = TRUE, estimate_prior_variance = TRUE, verbose = TRUE, min_abs_cor = 0.1*.

We binned all nfrQTL and nucQTL 95% credible variants by their posterior inclusion probabilities (PIP). We defined PIP bins ranging between 0 to 1 in increments of 0.1. For each bin, we calculated the fraction of nfrQTL or nucQTL credible variants overlapped their corresponding nfrPeak or nucPeak (*Fraction = Number of credible variants in PIP range and overlapping peak / Total number of credible variants in PIP range*).

We used coloc v5^8^ to test for colocalization between pairs of nfrQTLs and nucQTLs with summit-to-summit distance <5kb. We only considered nfrQTLs and nucQTLs that had at least one 95% credible set. We considered nfrQTLs and nucQTLs as colocalized if the posterior probability of sharing the same causal variant (PPH4) was greater than 0.8. Density plots were generated, and local maxima were identified using the *find_peaks* function from the *scipy.signal* package, which detects peak positions along the density curve.

### Assessing association between nucQTL colocalization and eQTL overlap

To access whether an nfrQTL that is an eQTL is more likely to have a colocalized nucQTL than an nfrQTL with no evidence of eQTL effects, we fitted a logistic regression model, adjusting for the mean accessibility score of the nfrQTL region. The logistic regression model was implemented in R using the ‘glm’ function as follows: *glm(whether_eqtl ∼ nuc_coloc + mean_nfr, family = binomial(link = “logit”))*.

### Causal inference testing

We inferred the causal directionalities for all pairs of colocalized nfr and nucQTLs (PPH4 > 0.8) and ran the mediation analysis through the causal inference test (CIT, 2.3.10)^35^. For each pair of nfrQTL and nucQTL tested, we treated the lead variants inferred by the SUSiE model as the genetic instrumental variable (L), and we treated each peak’s normalized count or score as the effect in the model (G/T). We included relevant covariates, including, age, sex, BMI, batch, number of nuclei per cluster, TSS enrichment, and sample’s median fragment length in chromosome 22. CIT is a regression-based method, and it assumes that if a variant (L) influences an outcome (T) exclusively through an exposure (G), the outcome should be independent of the variant when conditioned on the exposure. We considered four models through the *cit.cp* function i) nfrQTL variant -> nfrPeak -> nucPeak (P nfr-nuc-causal), ii) nfrQTL variant -> nucPeak -> nfrPeak (P nfr-nuc-rev-causal), iii) nucQTL variant -> nfrPeak -> nucPeak (P nuc-nfr-rev-causal), and iv) nucQTL variant -> nucPeak -> nfrPeak (P nuc-nfr-causal). We ran each model with 100 permutations and calculated the FDR using *fdr.cit.* We considered a colocalized QTL pair to have nfr-to-nuc causal directionality if and only if FDR nfr-nuc-causal < 0.05, FDR nfr-nuc-rev-causal > 0.05, FDR nuc-nfr-rev-causal < 0.05, and FDR nuc-nfr-causal > 0.05. The nuc-to-nfr directionality was determined by inverse criteria.

### Motif enrichment analyses

We conducted several motif enrichment analyses using the MEME suite’s SEA tool (version 5.5.5)^49^. Our initial analyses used the HOCOMOCO v12 motif database’s 949 most common motifs that scored the best across their bendmarking datasets^50^. We began by checking the enrichment in previously defined nfrPeaks, nucPeaks, and caPeaks, generating sequences based on the hg38 reference file. Analyses were conducted with second order shuffled input sequences as controls (*seed=12*). For each motif and cell type, we computed an enrichment ratio (ENR_RATIO; foreground/background occurrence).

When comparing the enrichment difference at the peaks level, we focused on 143 motifs and those with ENR_RATIO > 1.5 in at least one category. Those 143 motifs include lineage-defining regulators for skeletal muscle (MEF2A–D, MYF5/6, PAX7, SIX1–6), pancreatic islet/endoderm (FOXA1–3, HNF1A/B, HNF4A/B, PAX4/6, NKX2.x/3.x/6.x), developmental homeobox classes (HOXA cluster, PAX1–9, PITX1–3, OTX1–2, CDX1/2/4, DMBX1, GSC/GSC2, CRX), nuclear receptor and metabolic sensors (RXRA, VDR, HNF4A/B), ETS/co-regulators and architectural factors (ELF1–5, FLI1, RFX1–6, YY1/YY2), GC-rich generalist families (SP1–5/7–9; KLF1–17), and immune/stress-responsive TFs (ATF1–4/6/6B/7, NFKB1/2, BCL6/BCL6B). Paralogous motifs were retained within families where subtle motif preferences can be informative (e.g., MEF2, SIX, RFX, SP, KLF, HOXA, PAX, NKX).

Following the same methodology, we also performed enrichment analysis on sequences with significant nfrQTLs or nucQTLs. To investigate motifs enriched in colocalized nfrQTLs, we selected nfrPeak sequences near colocalized nucQTLs with a posterior probability (PPH4) greater than 0.8. We used nfrQTL peaks without colocalized nucQTLs (PPH4 < 0.2) for controls. Motifs were retained for visualization if ENR_RATIO > 5 in at least one cell type.

Similarly, to check on motifs in nfrPeaks with nfr-to-nuc directionality, we used those nfrPeaks sequences as input and the second order shuffled sequences as control. Motifs were kept for visualization if ENR_RATIO > 3 and if significant (FDR < 0.05) in at least one cell type. Endothelial and FAP cells were excluded for visualization due to low power (not enough peaks with nfr-to-nuc directionality).

### Motif disruption analyses

We used MotifbreakR^51^ to explore how transcription factor motifs were altered by candidate variants. We built a custom motif library from HOCOMOCO v12, ran the analysis under *threshold = 0.01, method = “default”, bkg = c(A=0.25, C=0.25, G=0.25, T=0.25)*, and filtered the results by p<0.01. We focused on the motifs with >75% sequence match on the allele that increased accessibility.

### GWAS enrichment analyses

We assessed the enrichment of various GWAS variants in ATAC-seq peaks with significant nfrQTLs, including 384 curated GWAS traits as previously described^19^. Summary statistics were formatted and converted to hg38 coordinates. Stratified linkage disequilibrium score regression (LDSC) was used to test enrichment for each trait in cell type-specific ATAC-seq peaks with significant nfrQTLs. FDR q-values were calculated using the Benjamini-Hochberg method. Seven traits of interest, including T2D^37^, basal metabolic rate (UKBB, 23105^36^), fasting insulin^38^, creatinine (UKBB, 30700^36^), atrial fibrillation^39^, standing height (UKBB, 50^36^), and left hand grip strength (UKBB, 46^36^) were selected for fine mapping based on FDR and relevance to muscle tissue.

### nfrQTL and GWAS colocalization analysis

To create fine-mapped credible sets, we identified lead variants for selected GWAS summary statistics. For traits like T2D, we included lead variants reported^37^. For other traits without reported lead variants, we developed custom scripts to identify lead variants in closely flanking regions (±150kb) with p-values < 5 × 10^−8^. For traits not in GRCh38, we lifted the variants from GRCh37 to GRCh38. For traits that include summary statistics for multiple ancestries, we only used the effect sizes and standard errors of European ancestry statistics for fine mapping. We also checked the reference alleles’ alignment between the GWAS summary statistics and our samples’ genotype information. For those variants that did not align, we reversed the effect size to *\*-1*. Fine-mapped credible sets were constructed around these lead variants using the Sum of SIngle Effects (SuSiE) model^3^, with settings including *L = 10, estimate_residual_variance = TRUE, estimate_prior_variance = TRUE, verbose = TRUE, and min_abs_cor = 0.1*. For locations where credible sets could not be formed within these parameters, we proceeded using effect sizes, variance of effect sizes, and the LD matrix to build credible set.

For GWAS lead variants where a 95% credible set could be formed and were in close proximity (within 100kb) to an nfrQTL credible set, colocalization analysis was conducted using coloc.v5 ^8^. A GWAS signal and nfrQTL were identified as colocalized if the posterior probability of them sharing the same causal variant (PPH4) exceeded 0.5. We also referred back to the nfrQTL’s nominal pass scan results to prioritize variants that lie within the nfrPeaks for further evaluation.

## Data and code availability

Raw muscle snATAC-seq sequencing, participant genotyping, and participant phenotypes are available through dbGaP (phs001048.v3.p1). The nfrPeak and nucPeak count matrices and full nfrQTL and nucQTL summary statistics are available on Zenodo (https://doi.org/10.5281/zenodo.15620988). Complete code used to generate results in this manuscript are available at https://github.com/ParkerLab/muscle_nfr_nucQTL. The curated set of 384 GWAS summary statistics used in **Fig 5a** and **Supplementary Fig 12** were obtained as previously described^19^. Summary statistics for the seven muscle-related traits used in **Fig 5b**, **Supplementary Fig 13**, and **Table 1** were obtained through the cited references as follows: grip strength (UKBB, 46^36^), standing height (UKBB, 50^36^), basal metabolic rate (UKBB, 23105^36^), creatinine (UKBB, 30700^36^), type 2 diabetes (T2D)^37^, fasting insulin^38^, and atrial fibrillation^39^.

## Acknowledgements

We would like to thank the Foundation of National Institutes of Health (FNIH) and the Accelerated Medicines Partnerships Type 2 Diabetes (AMP T2D) initiative for support. This work was supported by grants R01DK117960, R01DK062370, R01DK129469, UM1DK126185, and P30AR069620. CCR was supported by DK007245.

## Author contributions

SCJP designed the study. XW performed computation analyses under supervision of CCR and SCJP. AV and PO provided additional bioinformatic support and guidance. ML, JT, TAL, KLM, MB, LJS, HAK, and FSC provided sample data through the FUSION study. KM, MB, FSC, LJS and SCJP provided supervision and mentoring. XW and CCR wrote the manuscript under supervision of SCJP. All authors reviewed and approved the manuscript.

---

**Supplementary Figure 1.**
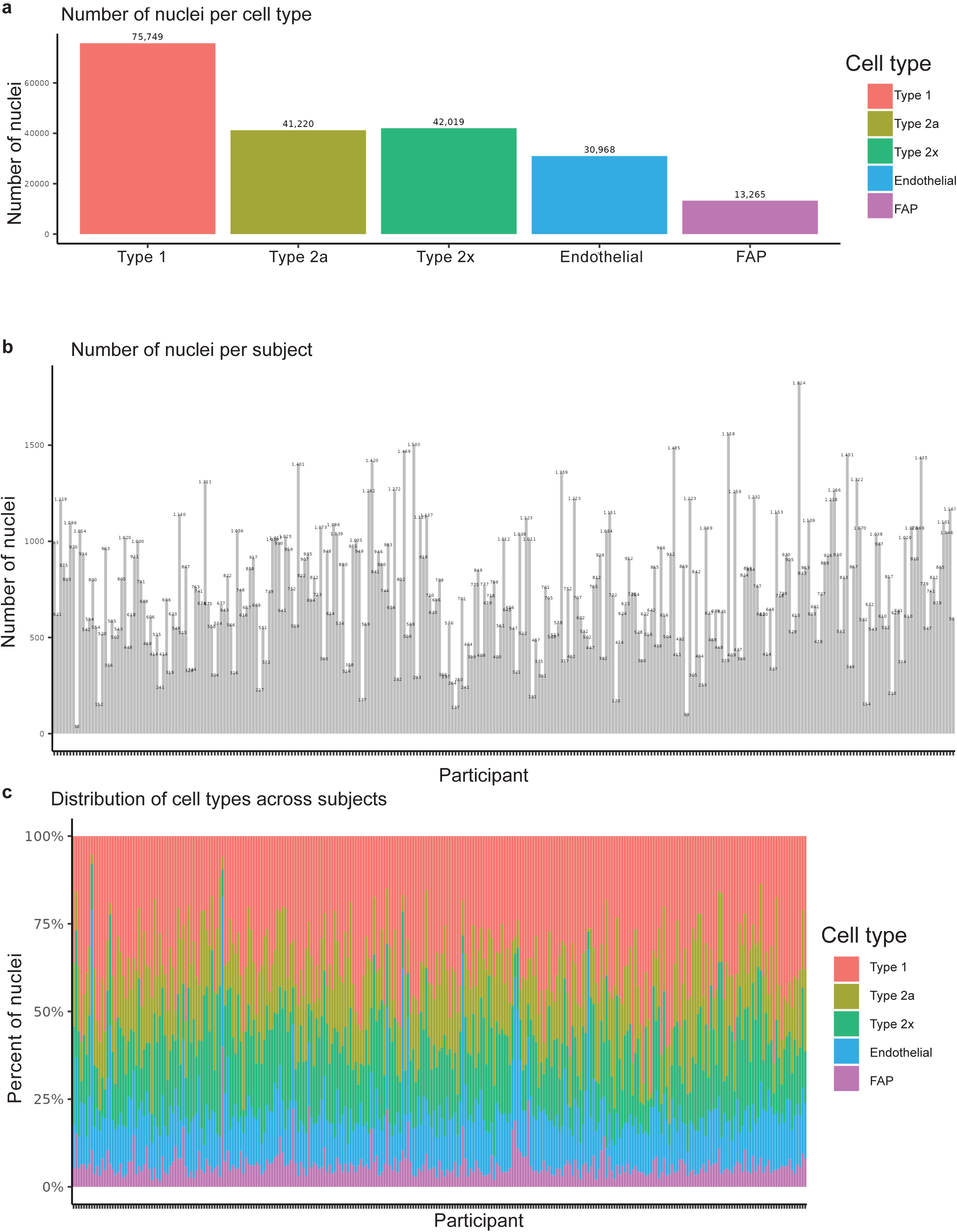
Overview of skeletal muscle snATAC-seq data used in this study. **a**. Number of nuclei called per cell type. **b**. Number of nuclei per participant. **c**. Distribution of the proportion of each cell type across participants.

**Supplementary Figure 2.**
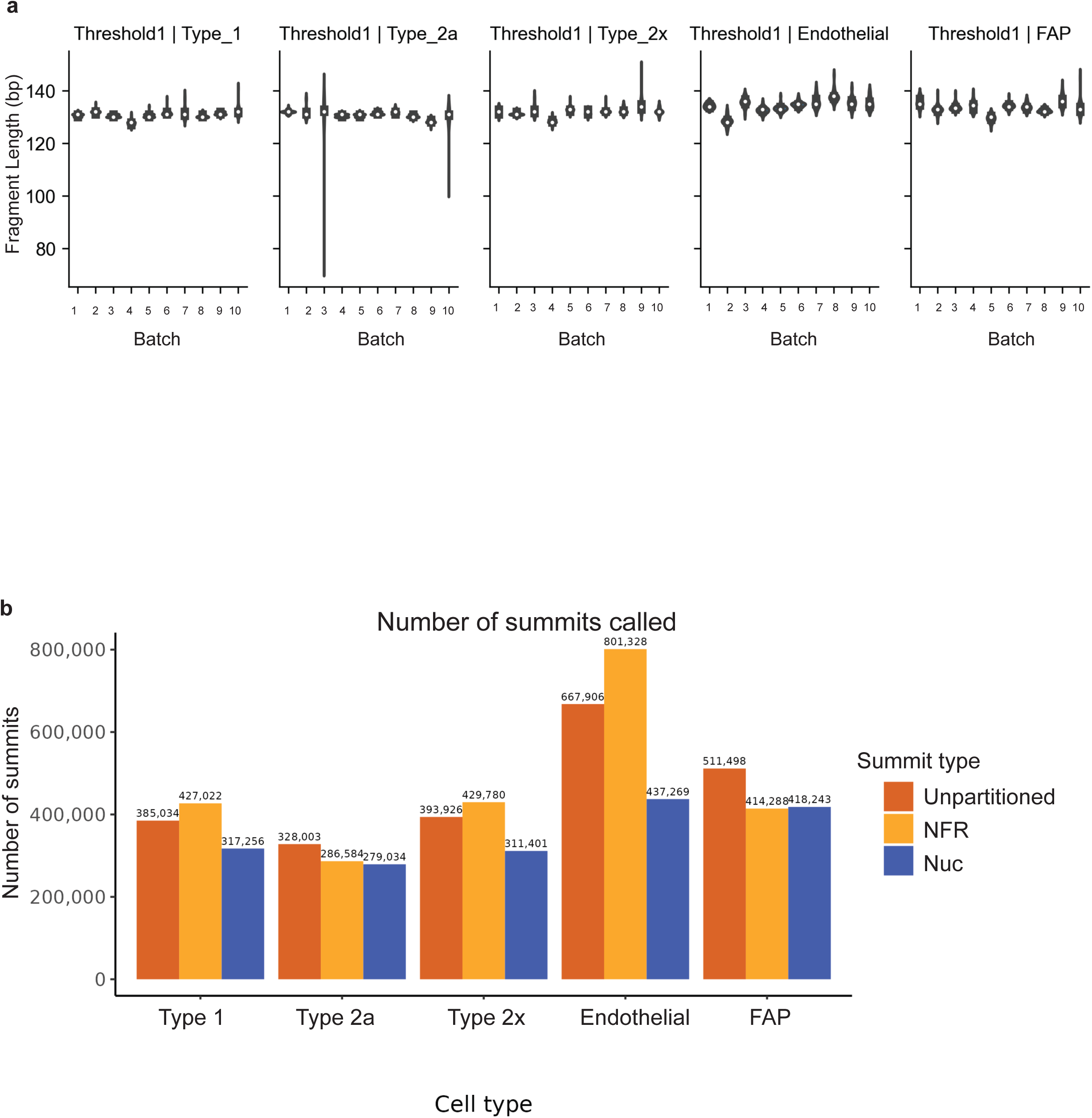
Partitioning ATAC-seq fragments and calling summits. **a**. Per-sample fragment length thresholds by batch and cell type as determined by multi-Otsu thresholding. **b**. The number of summits called by summit type and cell type.

**Supplementary Figure 3.**
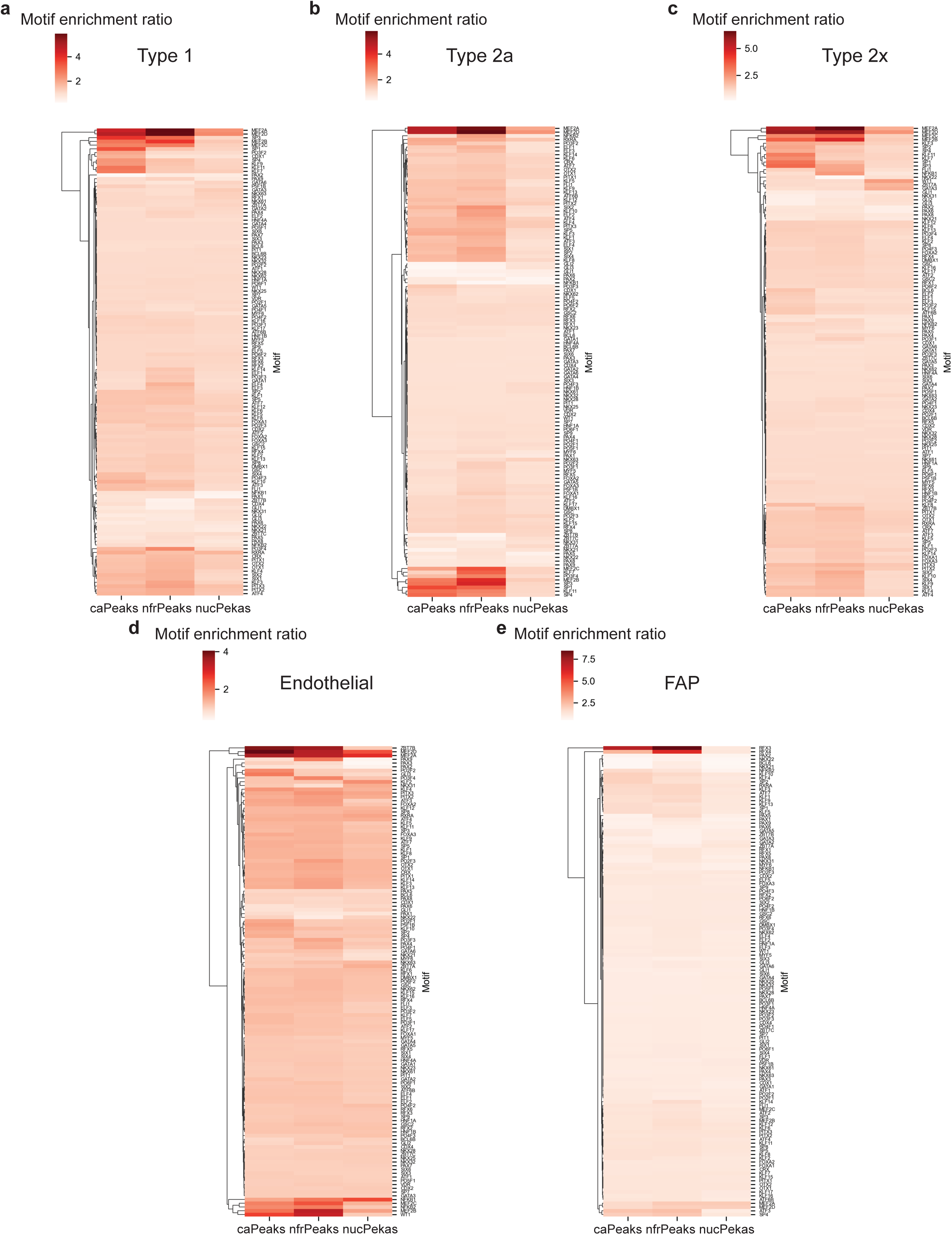
Motif enrichment analyses conducted on caPeaks, nfrPeaks, and nucPeaks using the HOCOMOCO v12 database. Enrichment ratios of selected motifs (n=125) are presented for **a**. type 1, **b**. type 2a, **c**. type 2x, **d**. endothelial, and **e**. FAP cells.

**Supplementary Figure 4.**
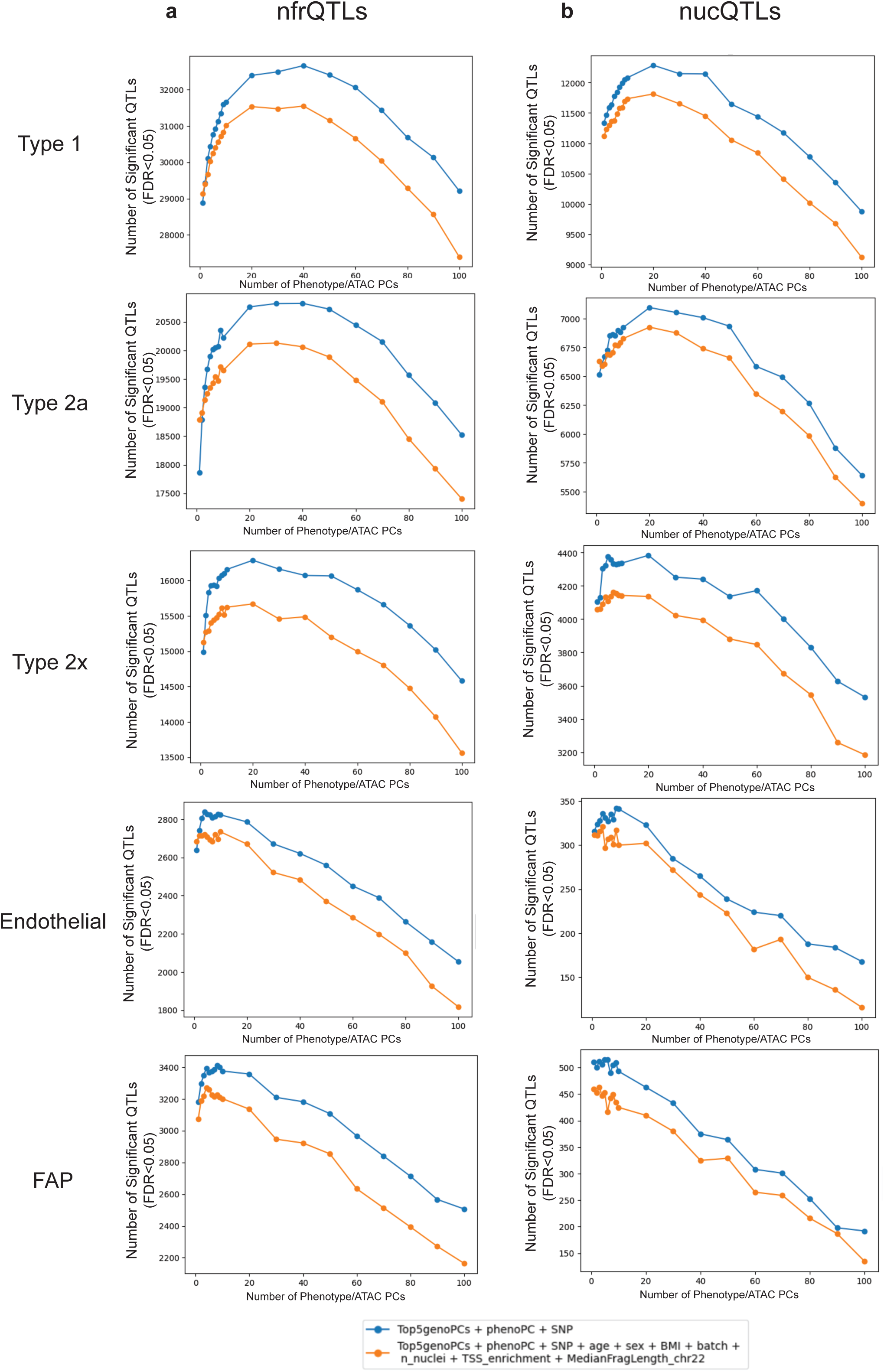
The number of peaks of with significant QTLs (FDR <5%). **a**. nfrQTLs and **b**. nucQTLs with models including explicit technical covariates (orange) or latent covariates only (blue).

**Supplementary Figure 5.**
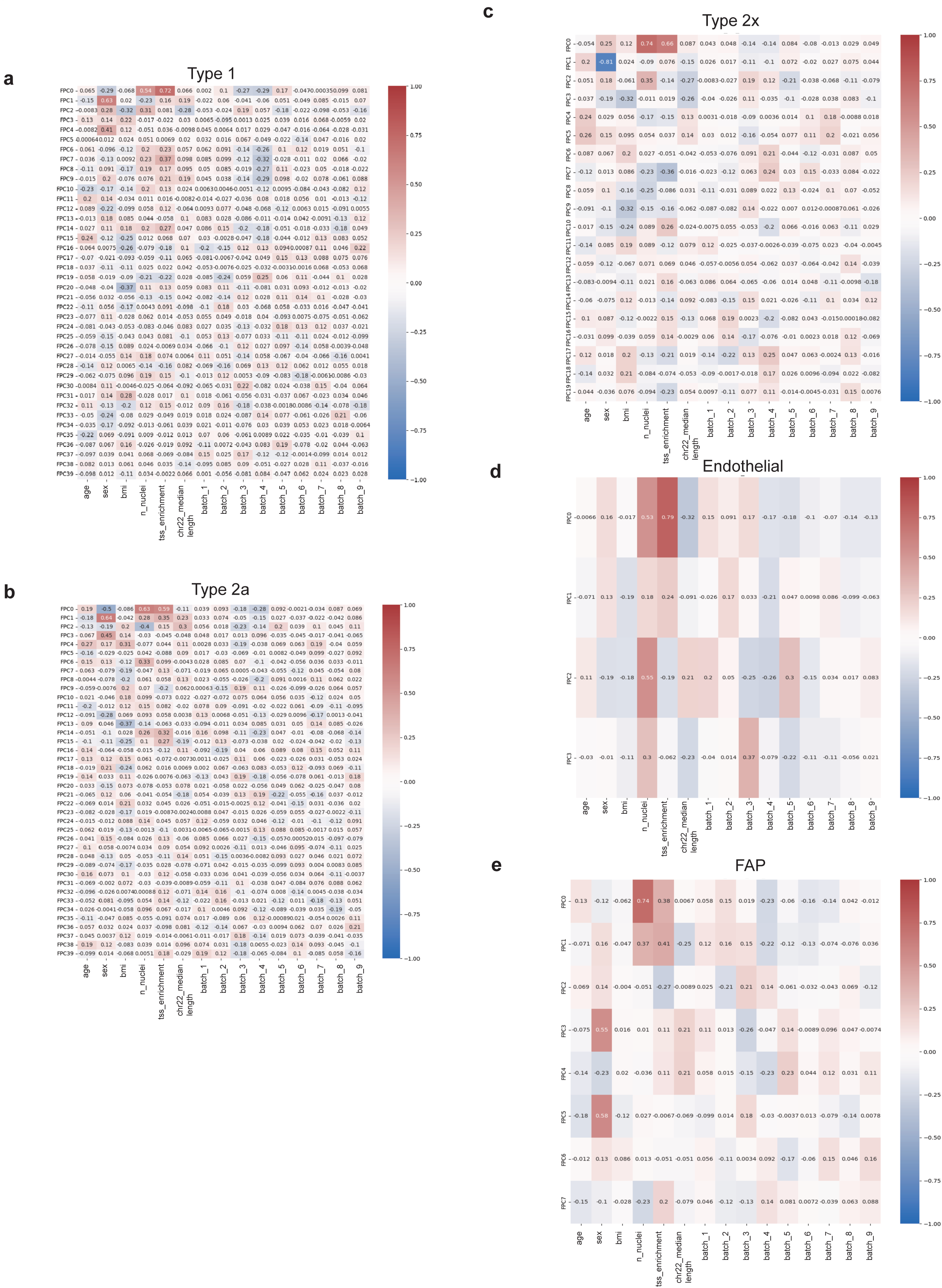
Correlation between phenotype PCs (FPCs) and technical covariates. **a.** type 1, **b.** type 2a, **c.** type 2x, **d.** endothelial, and **e.** FAP nfr regions.

**Supplementary Figure 6.**
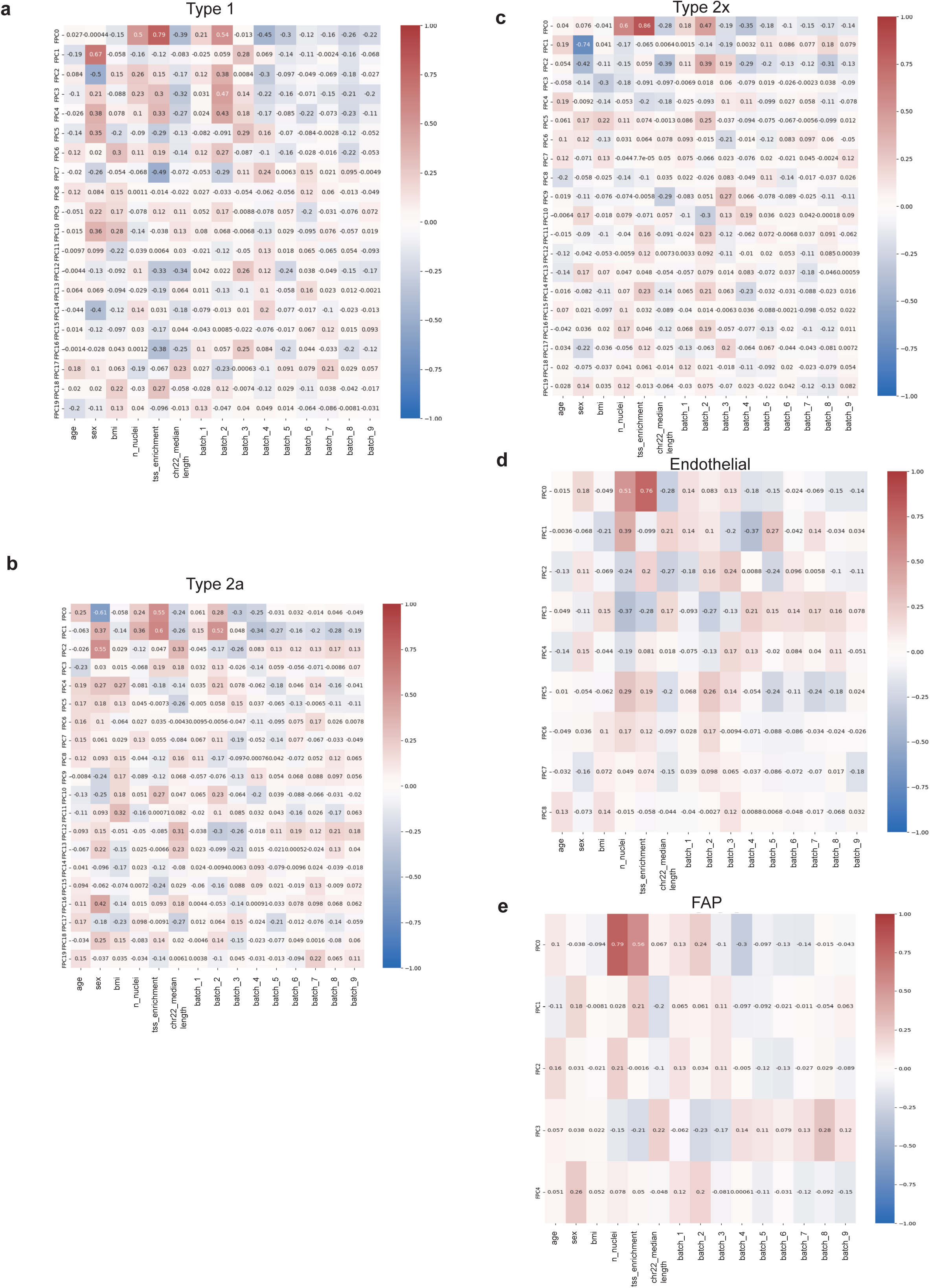
Correlation between phenotype PCs (FPCs) and technical covariates. **a.** type 1, **b.** type 2a, **c.** type 2x, **d.** endothelial, and **e.** FAP nuc regions.

**Supplementary Figure 7.**
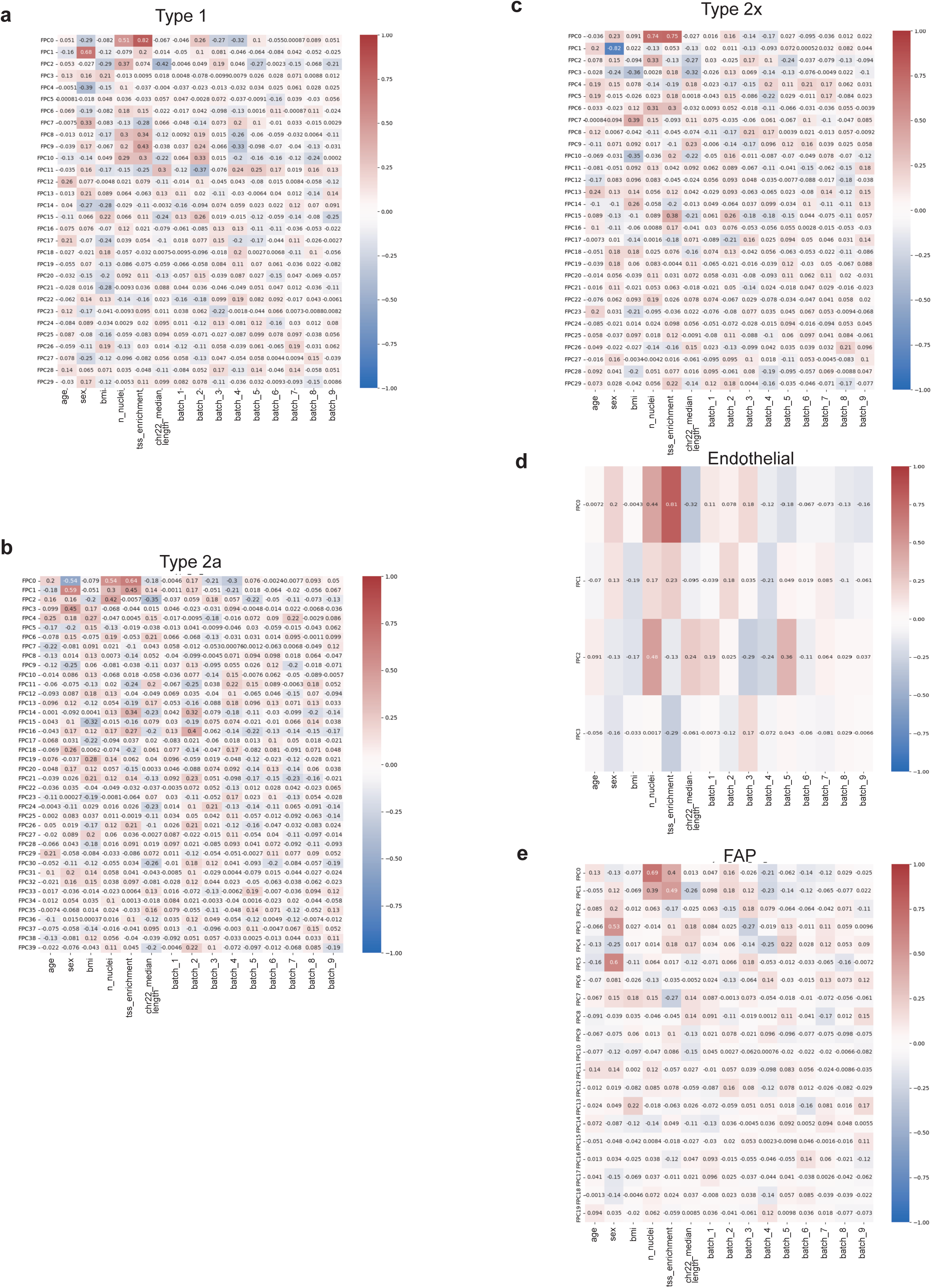
Correlation between phenotype PCs (FPCs) and technical covariates. **a.** type 1, **b.** type 2a, **c.** type 2x, **d.** endothelial, and **e.** FAP ca regions.

**Supplementary Figure 8.**
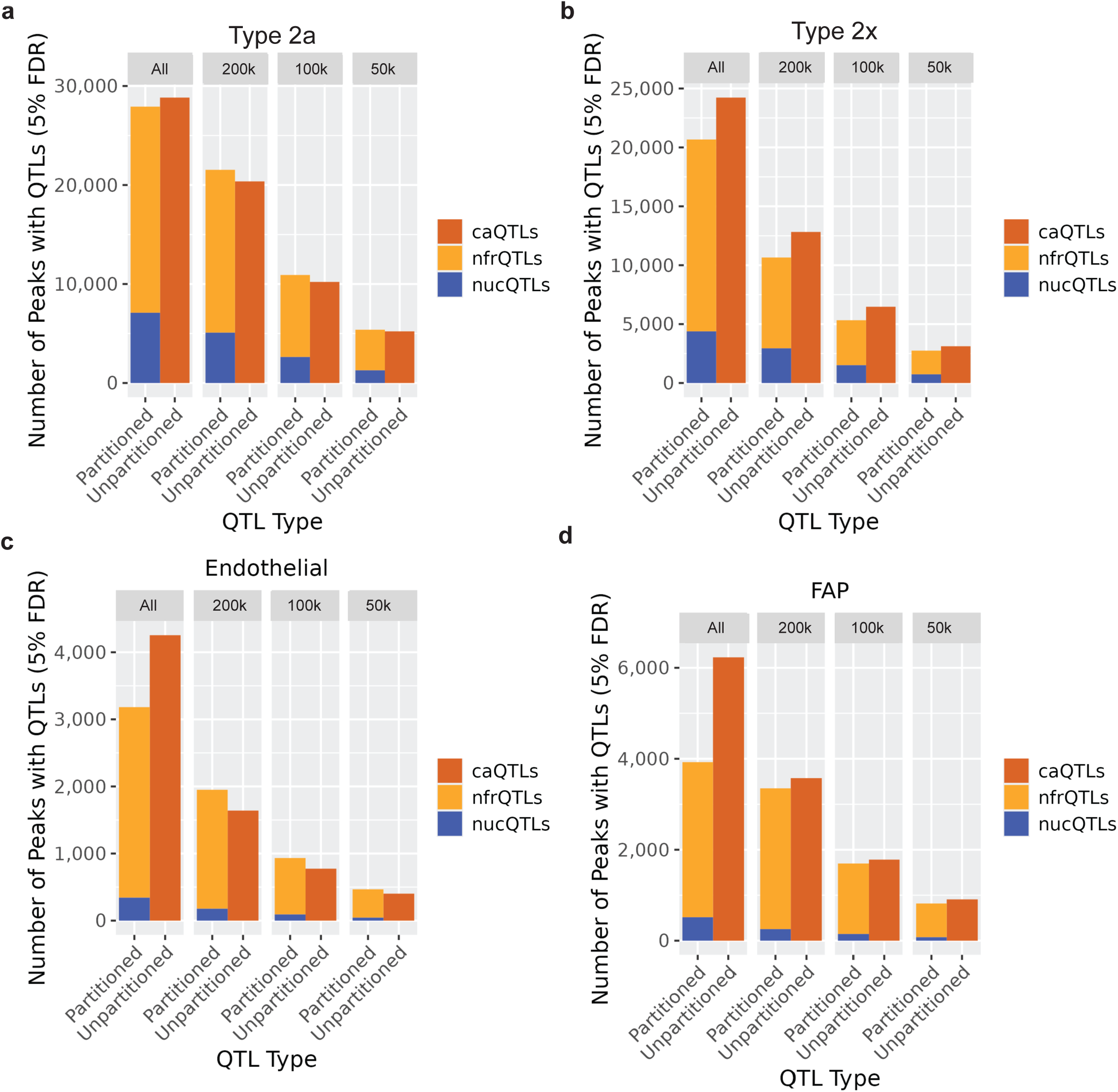
The number of peaks with significant QTLs (FDR<5%) stratified by the number of peaks included at the start of the QTL scan. **a**. type 2a, **b**. type 2x, **c**. endothelial, and **d**.FAP cells. Figures are faceted by the total number of peaks included in the QTL scan, including either all peaks (no downsampling) or downsampling to 200k, 100k, or 50k peaks.

**Supplementary Figure 9.**
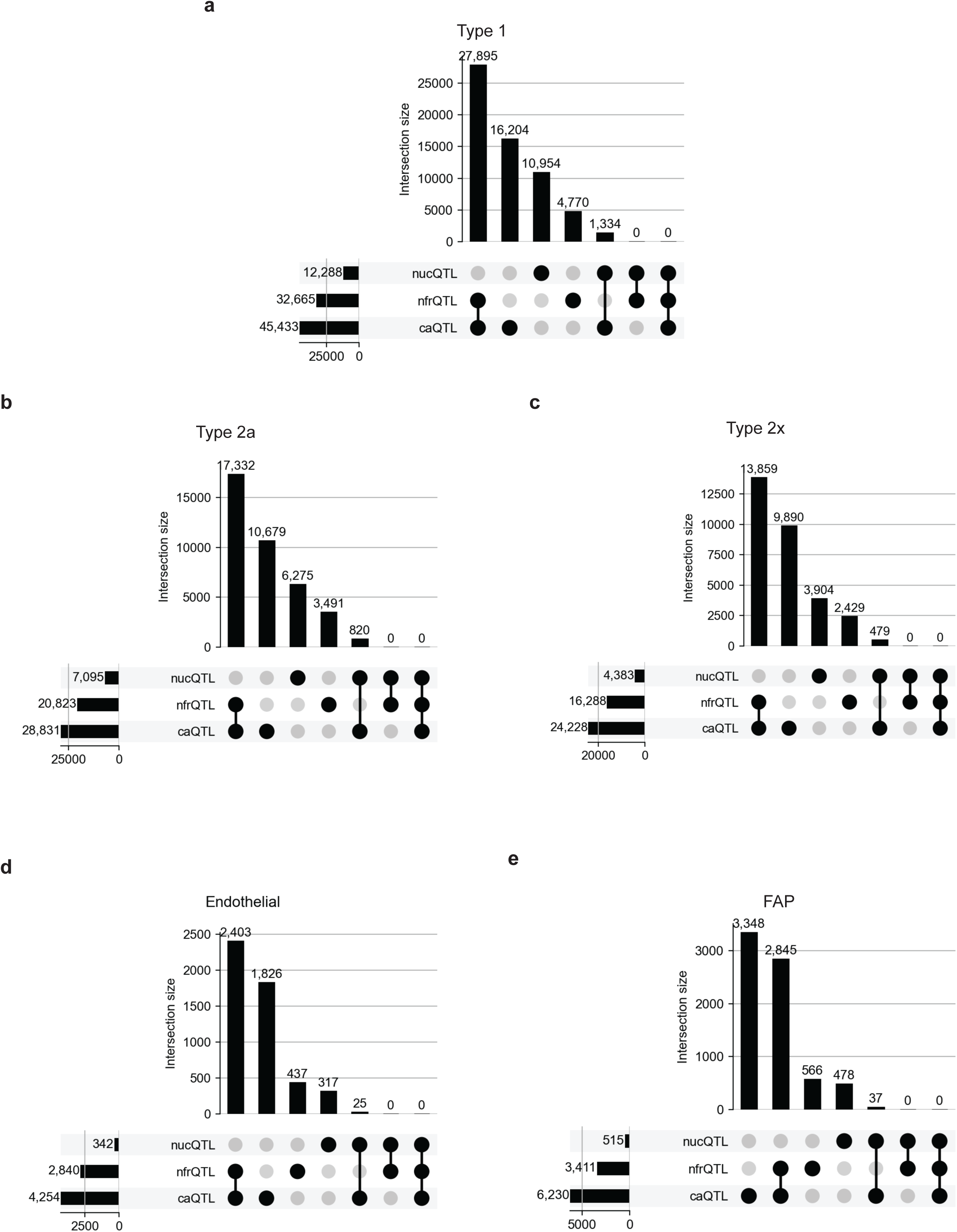
Upset plots depicting QTL peak overlap, where two peaks were considered to overlap if at least 50% of both peaks overlapped the other peak. **a**. type 1, **b**. type 2a, **c**. type 2x, **d**. endothelial, and **e**. FAP cells.

**Supplementary Figure 10.**
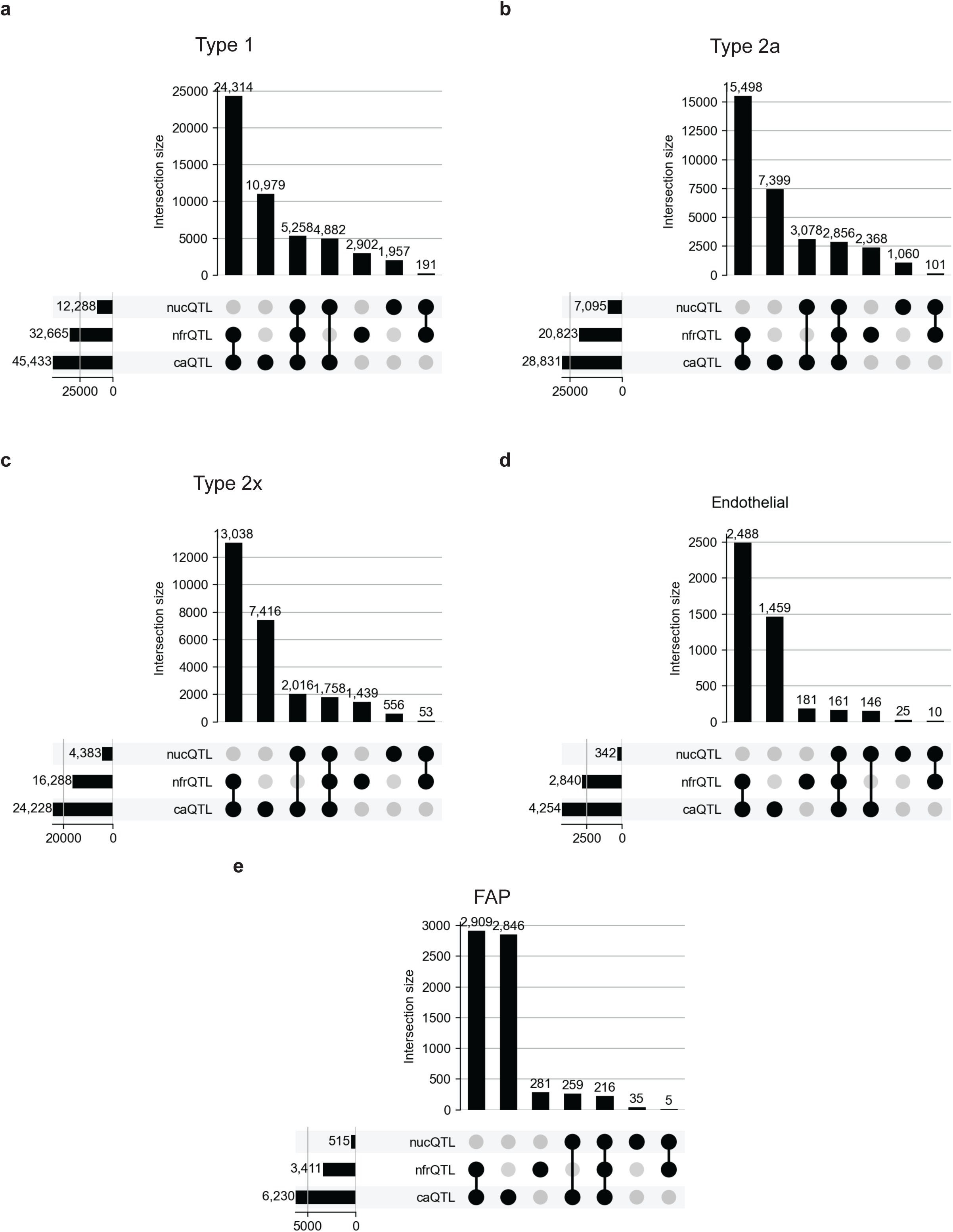
Upset plots depicting QTL peak overlap, where two peaks were considered to overlap if at least 1bp is overlapping. **a**. type 1, **b**. type 2a, **c**. type 2x, **d**. endothelial, and **e**. FAP cells.

**Supplementary Figure 11.**
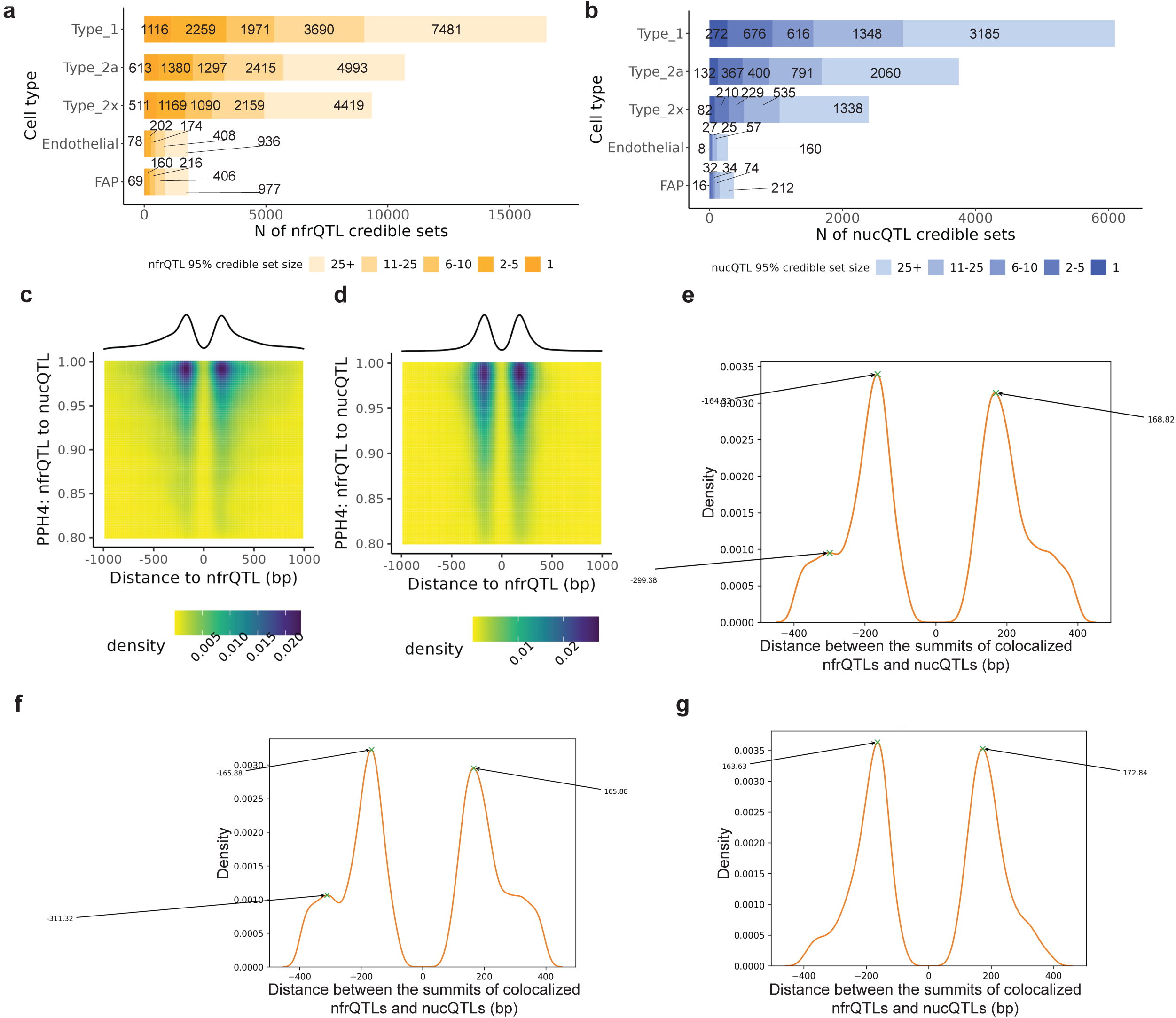
Fine mapping and colocalization for nfrQTLs and nucQTLs. The number of **a**. nfrQTL and **b**. nucQTL credible sets detected by fine mapping for each cell type by credible set size. **c-d**. Relationship between evidence for colocalization and distance between nucQTLs and nfrQTLs. **e-g**. Estimating local maxima from density plots to determine typical distance between colocalizing nucQTL and nfrQTL pairs. **c & f**. Pairs where the nfrQTL colocalizes with more than one nucQTL (n=13,379). **d & g**. Pairs where the nfrQTL colocalized with only one nucQTL (n=3,539). **e**. All colocalized pairs (n=16,918) (corresponds to heatmap and density from Fig 3a).

**Supplementary Figure 12.**
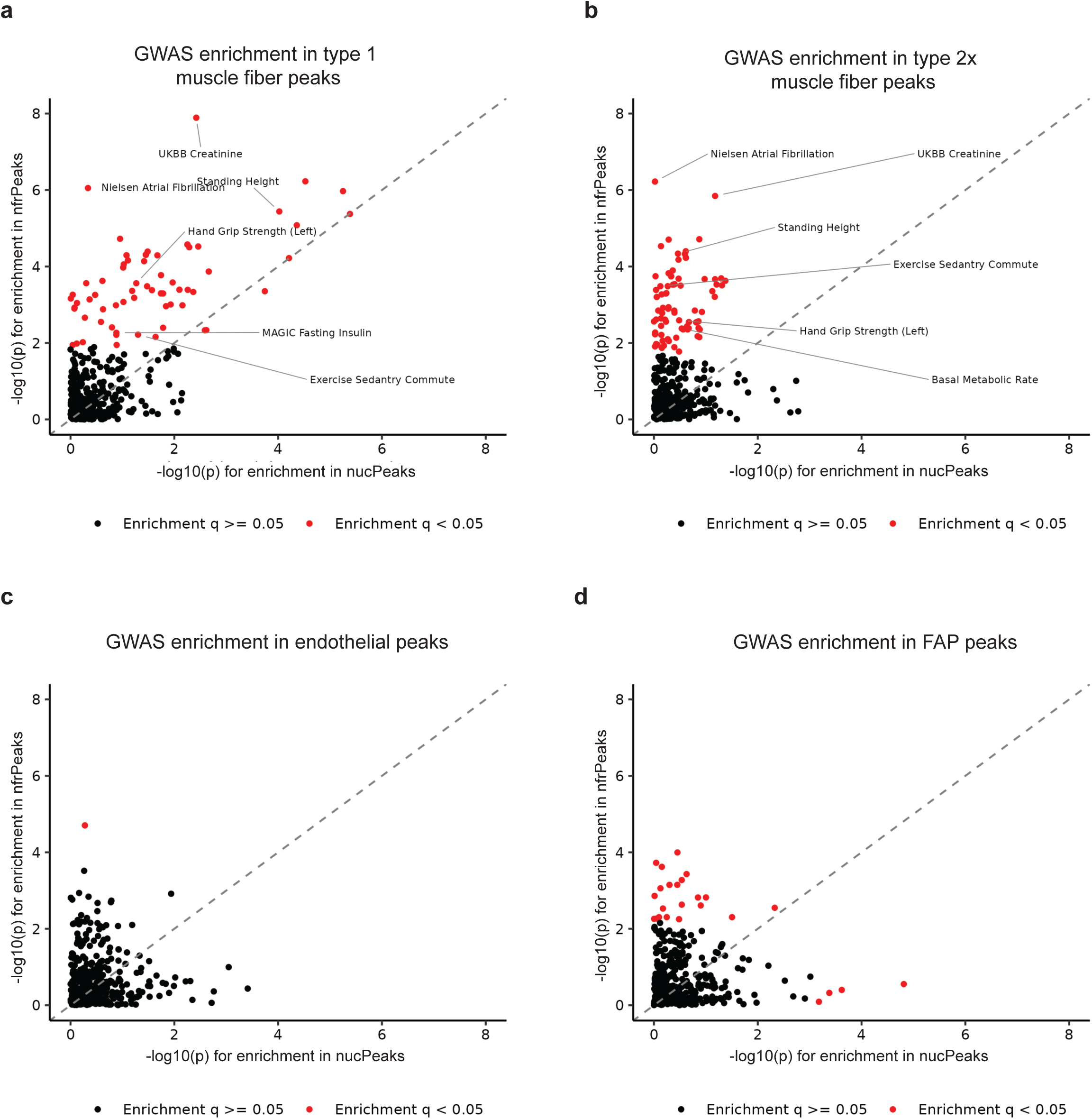
GWAS enrichment in nfrQTL and nucQTL regions. Each point represents a GWAS trait (n=384 traits). LDSC -log10(p-value) for enrichment in nucQTL regions (x-axis) and nfrQTL regions (y-axis) for **a**. type 1, **b**. type 2x, **c**. endothelial, and **d**. FAP cells. Traits with significant enrichment (FDR<5%) in either region set are colored red. LDSC = Linkage disequilibrium score regression.

**Supplementary Figure 13.**
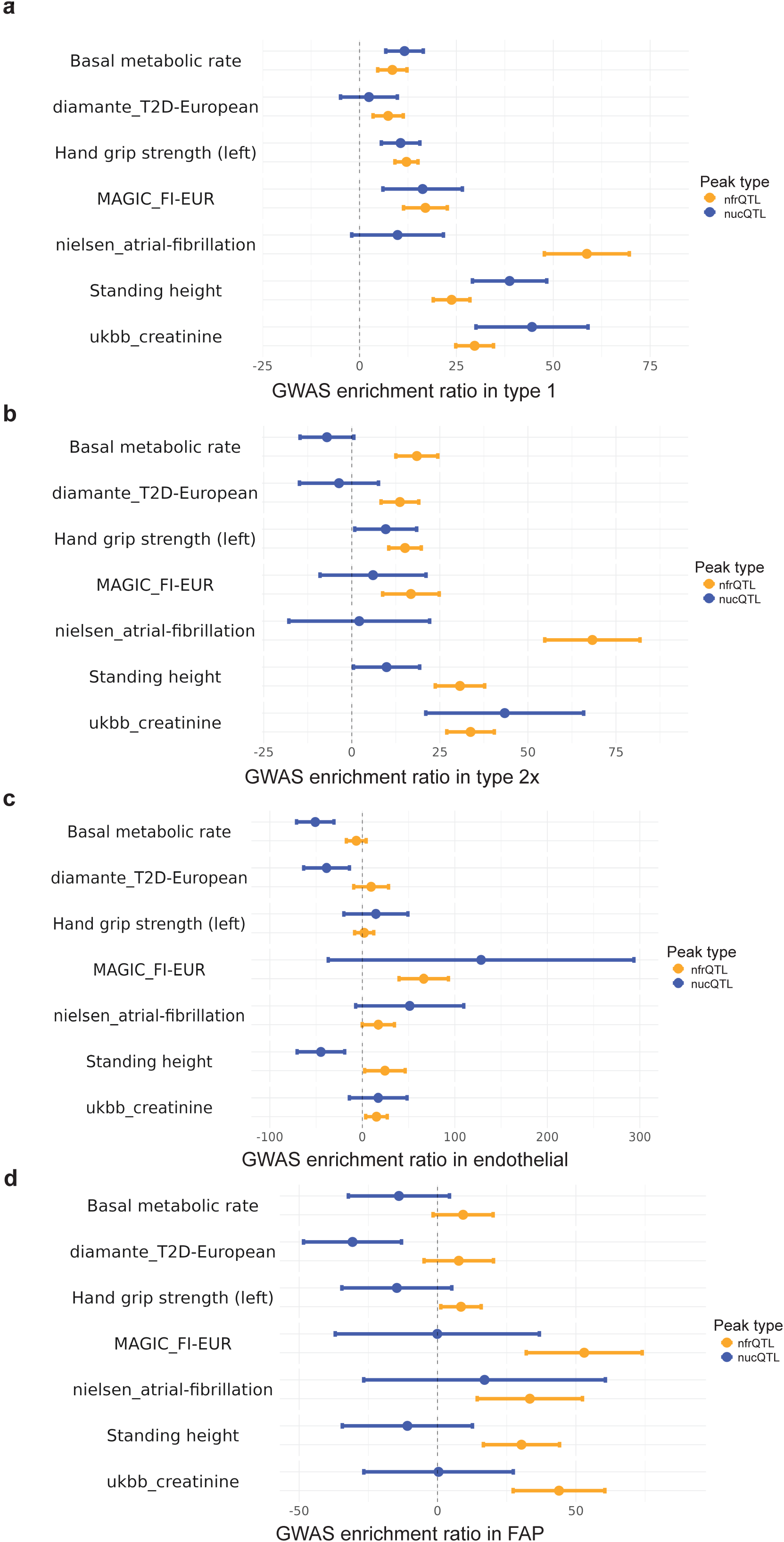
Forest plots for GWAS enrichment in nfrQTL and nucQTL regions. **a.** type 1, **b**. type 2x, **c**. endothelial, and **d**. FAP cells. Bars show estimated enrichment and standard error.

**Supplementary Figure 14.**
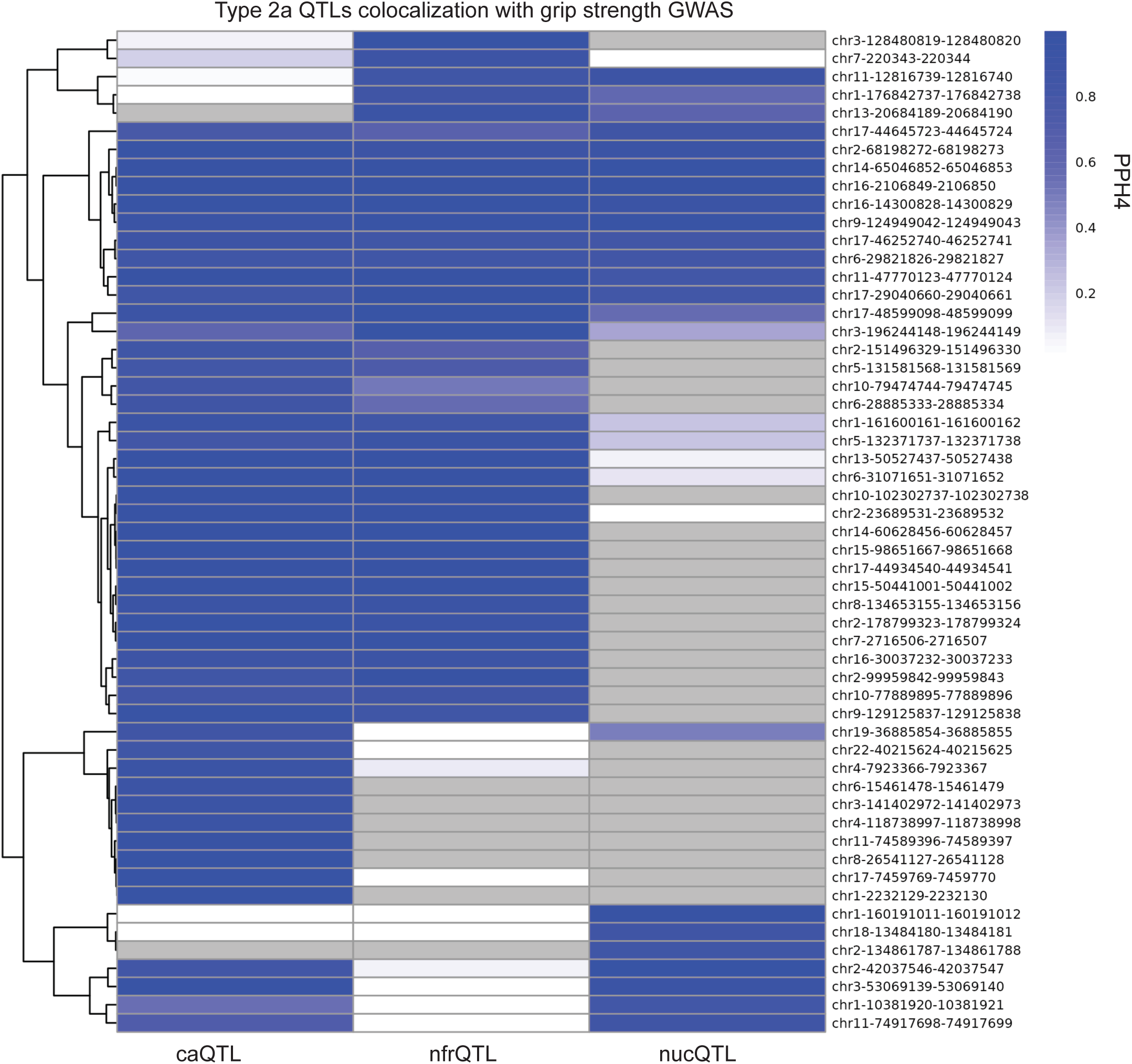
Colocalization of grip strength GWAS signals with caQTLs, nfrQTLs, and nucQTLs in type 2a muscle fiber.

**Supplementary Table 1.**
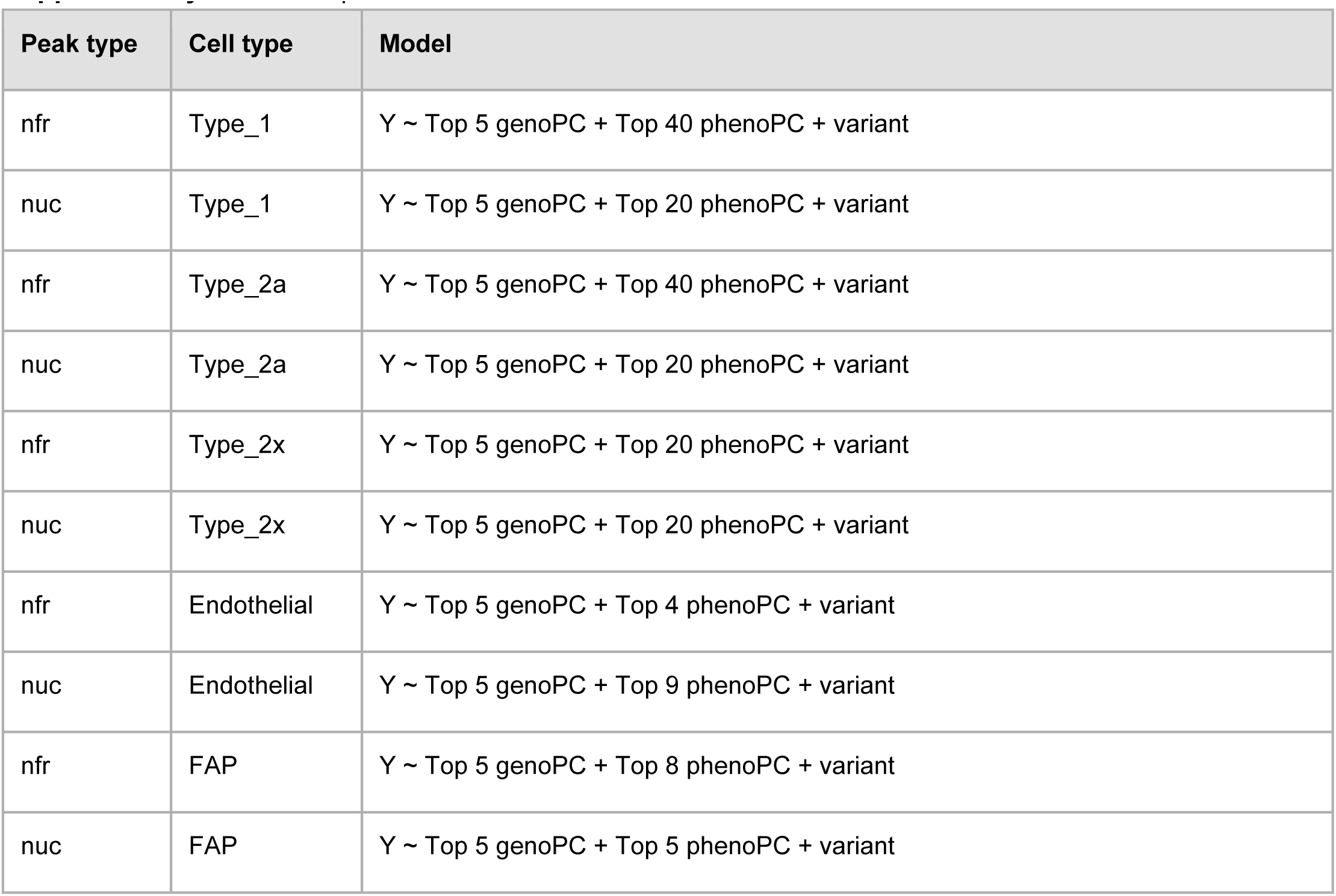
Optimized statistical models used for *cis-*QTL scans.

| Peak type | Cell type | Model |
| --- | --- | --- |
| nfr | Type_1 | Y ~ Top 5 genoPC + Top 40 phenoPC + variant |
| nuc | Type_1 | Y ~ Top 5 genoPC + Top 20 phenoPC + variant |
| nfr | Type_2a | Y ~ Top 5 genoPC + Top 40 phenoPC + variant |
| nuc | Type_2a | Y ~ Top 5 genoPC + Top 20 phenoPC + variant |
| nfr | Type_2x | Y ~ Top 5 genoPC + Top 20 phenoPC + variant |
| nuc | Type_2x | Y ~ Top 5 genoPC + Top 20 phenoPC + variant |
| nfr | Endothelial | Y ~ Top 5 genoPC + Top 4 phenoPC + variant |
| nuc | Endothelial | Y ~ Top 5 genoPC + Top 9 phenoPC + variant |
| nfr | FAP | Y ~ Top 5 genoPC + Top 8 phenoPC + variant |
| nuc | FAP | Y ~ Top 5 genoPC + Top 5 phenoPC + variant |

**Supplementary Table 2.** Number of PCs required to explain 90% of the variance in accessibility profiles among peaks with significant QTLs.

| Cell type | Number of PCs explaining 90% of caQTL peak variance | Number of PCs explaining 90% of nfrQTL and nucQTL peak variance |
| --- | --- | --- |
| Type_1 | 205 | 210 |
| Type_2a | 196 | 209 |
| Type_2x | 204 | 218 |
| Endothelial | 224 | 224 |
| FAP | 210 | 221 |

**Supplementary Table 3.** Summary of colocalization between nfrQTLs and nucQTLs.

| Cell type | Number of pairs studied | Number of coloc pairs (PPH4 > 0.5) | Number of coloc pairs (PPH4 > 0.8) | Number of unique nfrQTLs in pairs (PPH4 > 0.8) | Number of nfrQTLs colocalized with >1 nucQTLs (PPH4 > 0.8) | Number of unique nucQTLs in pairs (PPH4 > 0.8) | Number of nucQTLs colocalized with >1 nfrQTLs (PPH4 > 0.8) |
| --- | --- | --- | --- | --- | --- | --- | --- |
| Type_1 | 12,855 | 9,849 | 8,473 | 4,305 | 2,162 | 4,525 | 2,103 |
| Type_2a | 7,089 | 5,605 | 4,781 | 2,485 | 1,186 | 2,710 | 1,160 |
| Type_2x | 4,431 | 3,589 | 3,036 | 1,717 | 730 | 1,689 | 736 |
| Endothelial | 436 | 344 | 294 | 178 | 60 | 179 | 74 |
| FAP | 455 | 391 | 334 | 225 | 75 | 222 | 77 |
| Total | 25,266 | 19,778 | 16,918 | 8,910 | 4,213 | 9,325 | 4,150 |

## References

1. Cerezo, M. et al. The NHGRI-EBI GWAS Catalog: standards for reusability, sustainability and diversity. Nucleic Acids Res. gkae1070 (2024) doi:10.1093/nar/gkae1070.

2. Maurano, M. T. et al. Systematic localization of common disease-associated variation in regulatory DNA. Science 337, 1190–1195 (2012).

3. Giambartolomei, C. et al. Bayesian Test for Colocalisation between Pairs of Genetic Association Studies Using Summary Statistics. PLOS Genet. 10, e1004383 (2014).

4. Gusev, A. et al. Integrative approaches for large-scale transcriptome-wide association studies. Nat. Genet. 48, 245–252 (2016).

5. Aguet, F. et al. Genetic effects on gene expression across human tissues. Nature 550, 204–213 (2017).

6. THE GTEX CONSORTIUM. The GTEx Consortium atlas of genetic regulatory effects across human tissues. Science 369, 1318–1330 (2020).

7. Viñuela, A. et al. Genetic variant effects on gene expression in human pancreatic islets and their implications for T2D. Nat. Commun. 11, 4912 (2020).

8. Wallace, C. A more accurate method for colocalisation analysis allowing for multiple causal variants. PLOS Genet. 17, e1009440 (2021).

9. Orchard, P. et al. Cross-cohort analysis of expression and splicing quantitative trait loci in TOPMed. 2025.02.19.25322561 Preprint at 10.1101/2025.02.19.25322561 (2025).

10. Umans, B. D., Battle, A. & Gilad, Y. Where Are the Disease-Associated eQTLs? Trends Genet. 37, 109–124 (2021).

11. GTEx Consortium. The GTEx Consortium atlas of genetic regulatory effects across human tissues. Science 369, 1318–1330 (2020).

12. Mostafavi, H., Spence, J. P., Naqvi, S. & Pritchard, J. K. Systematic differences in discovery of genetic effects on gene expression and complex traits. Nat. Genet. 55, 1866–1875 (2023).

13. Võsa, U. et al. Large-scale cis- and trans-eQTL analyses identify thousands of genetic loci and polygenic scores that regulate blood gene expression. Nat. Genet. 53, 1300–1310 (2021).

14. van der Wijst, M. G. P. et al. Single-cell RNA sequencing identifies cell type-specific cis-eQTLs and co-expression QTLs. Nat. Genet. 50, 493–497 (2018).

15. Natri, H. M. et al. Cell-type-specific and disease-associated expression quantitative trait loci in the human lung. Nat. Genet. 56, 595–604 (2024).

16. Bian, L. et al. Single-cell eQTL mapping reveals cell-type-specific genes associated with the risk of gastric cancer. Cell Genomics 5, 100812 (2025).

17. Arthur, T. D. et al. Multiomic QTL mapping reveals phenotypic complexity of GWAS loci and prioritizes putative causal variants. Cell Genomics 5, 100775 (2025).

18. Jeong, R. & Bulyk, M. L. Chromatin accessibility variation provides insights into missing regulation underlying immune-mediated diseases. eLife 13, (2024).

19. Varshney, A. et al. Population-scale skeletal muscle single-nucleus multi-omic profiling reveals extensive context specific genetic regulation. 2023.12.15.571696 Preprint at 10.1101/2023.12.15.571696 (2024).

20. Buenrostro, J. D., Wu, B., Chang, H. Y. & Greenleaf, W. J. ATAC-seq: A Method for Assaying Chromatin Accessibility Genome-Wide. Curr. Protoc. Mol. Biol. 109, 21.29.1–21.29.9 (2015).

21. Buenrostro, J. D. et al. Integrated Single-Cell Analysis Maps the Continuous Regulatory Landscape of Human Hematopoietic Differentiation. Cell 173, 1535–1548.e16 (2018).

22. Buenrostro, J. D., Giresi, P. G., Zaba, L. C., Chang, H. Y. & Greenleaf, W. J. Transposition of native chromatin for multimodal regulatory analysis and personal epigenomics. Nat. Methods 10, 1213–1218 (2013).

23. Preissl, S. et al. Single-nucleus analysis of accessible chromatin in developing mouse forebrain reveals cell-type-specific transcriptional regulation. Nat. Neurosci. 21, 432–439 (2018).

24. Zhang, Y. et al. Model-based Analysis of ChIP-Seq (MACS). Genome Biol. 9, R137 (2008).

25. Li, Z. et al. Identification of transcription factor binding sites using ATAC-seq. Genome Biol. 20, 45 (2019).

26. Baek, S., Goldstein, I. & Hager, G. L. Bivariate Genomic Footprinting Detects Changes in Transcription Factor Activity. Cell Rep. 19, 1710–1722 (2017).

27. D’Oliveira Albanus, R., et al. Chromatin information content landscapes inform transcription factor and DNA interactions. Nat. Commun. 12, 1307 (2021).

28. Dudek, M. F. et al. Characterization of non-coding variants associated with transcription-factor binding through ATAC-seq-defined footprint QTLs in liver. Am. J. Hum. Genet. 0, (2025).

29. Tarbell, E. D. & Liu, T. HMMRATAC: a Hidden Markov ModeleR for ATAC-seq. Nucleic Acids Res. 47, e91 (2019).

30. Schep, A. N. et al. Structured nucleosome fingerprints enable high-resolution mapping of chromatin architecture within regulatory regions. Genome Res. 25, 1757–1770 (2015).

31. Liao, P.-S., Chen, T.-S. & Chung, P.-C. A Fast Algorithm for Multilevel Thresholding. J. Inf. Sci. Eng. 17, 713–727 (2001).

32. Luger, K., Mäder, A. W., Richmond, R. K., Sargent, D. F. & Richmond, T. J. Crystal structure of the nucleosome core particle at 2.8 A resolution. Nature 389, 251–260 (1997).

33. Gómora-García, J. C. & Furlan-Magaril, M. Pioneer factors outline chromatin architecture. Curr. Opin. Cell Biol. 93, 102480 (2025).

34. Gryder, B. E. et al. PAX3–FOXO1 Establishes Myogenic Super Enhancers and Confers BET Bromodomain Vulnerability. Cancer Discov. 7, 884–899 (2017).

35. Millstein, J., Chen, G. K. & Breton, C. V. cit: hypothesis testing software for mediation analysis in genomic applications. Bioinformatics 32, 2364–2365 (2016).

36. UK Biobank. Neale lab http://www.nealelab.is/uk-biobank.

37. Mahajan, A. et al. Fine-mapping type 2 diabetes loci to single-variant resolution using high-density imputation and islet-specific epigenome maps. Nat. Genet. 50, 1505–1513 (2018).

38. Lagou, V. et al. Sex-dimorphic genetic effects and novel loci for fasting glucose and insulin variability. Nat. Commun. 12, 24 (2021).

39. Nielsen, J. B. et al. Biobank-driven genomic discovery yields new insight into atrial fibrillation biology. Nat. Genet. 50, 1234–1239 (2018).

40. Grand, R. S. et al. Genome access is transcription factor-specific and defined by nucleosome position. Mol. Cell (2024) doi:10.1016/j.molcel.2024.08.009.

41. Yi, C., Kitamura, Y., Maezawa, S., Namekawa, S. H. & Cairns, B. R. ZBTB16/PLZF regulates juvenile spermatogonial stem cell development through an extensive transcription factor poising network. Nat. Struct. Mol. Biol. 1–14 (2025) doi:10.1038/s41594-025-01509-5.

42. Gao, C. et al. Iterative single-cell multi-omic integration using online learning. Nat. Biotechnol. 39, 1000–1007 (2021).

43. Quinlan, A. R. & Hall, I. M. BEDTools: a flexible suite of utilities for comparing genomic features. Bioinformatics 26, 841–842 (2010).

44. Liao, Y., Smyth, G. K. & Shi, W. featureCounts: an efficient general purpose program for assigning sequence reads to genomic features. Bioinformatics 30, 923–930 (2014).

45. Ramírez, F. et al. deepTools2: a next generation web server for deep-sequencing data analysis. Nucleic Acids Res. 44, W160–165 (2016).

46. Amemiya, H. M., Kundaje, A. & Boyle, A. P. The ENCODE Blacklist: Identification of Problematic Regions of the Genome. Sci. Rep. 9, 9354 (2019).

47. Delaneau, O. et al. A complete tool set for molecular QTL discovery and analysis. Nat. Commun. 8, 15452 (2017).

48. Wang, G., Sarkar, A., Carbonetto, P. & Stephens, M. A Simple New Approach to Variable Selection in Regression, with Application to Genetic Fine Mapping. J. R. Stat. Soc. Ser. B Stat. Methodol. 82, 1273–1300 (2020).

49. Bailey, T. L. & Grant, C. E. SEA: Simple Enrichment Analysis of motifs. 2021.08.23.457422 Preprint at 10.1101/2021.08.23.457422 (2021).

50. Vorontsov, I. E. et al. HOCOMOCO in 2024: a rebuild of the curated collection of binding models for human and mouse transcription factors. Nucleic Acids Res. 52, D154–D163 (2024).

51. Coetzee, S. G., Coetzee, G. A. & Hazelett, D. J. *motifbreakR* : an R/Bioconductor package for predicting variant effects at transcription factor binding sites. Bioinformatics 31, 3847–3849 (2015).

